# Estrogen Receptors/E2F1/CDKN3 Axis Protects from UV-induced Skin Cancers in Females

**DOI:** 10.1101/2025.02.12.637819

**Authors:** Céline Lukowicz, Carine Winkler, Catherine Roger, Joanna C Fowler, Yi-Chien Tsai, Joachim Meuli, Stéphanie Claudinot, Yun-Tsan Chang, Christoph Iselin, Philip H Jones, Emmanuella Guenova, Paris Jafari, Liliane Michalik

## Abstract

Men have a significantly higher risk of developing cutaneous squamous cell carcinoma (SCC) compared to women, but models and comprehensive analysis of signaling pathways highlighting this sexual dimorphism are missing. In this study, we display a UV-induced SCC model in hairless mice recapitulating this sex difference, with enhanced SCC development in males. While UV-induced DNA damage is similar between sexes, we uncovered sex-specific responses in epidermal cell proliferation and differentiation. Using global transcriptional profile analyses, we identified E2F transcription factors as key sex-specific markers involved in the proliferative response to UV. Notably, E2F1/2, along with their target gene CDKN3, were selectively downregulated in female mice and human epidermis following UV exposure. Mechanistically, UV-induced and sex-specific modulation of E2F1 and CDKN3 expression is mediated by Estrogen Receptors. Lastly, low levels of CDKN3 in head and neck SCC are observed exclusively in female patients correlating with better prognosis. These findings shed new light on fundamental mechanisms protecting women from cancer after carcinogen exposure and could lead to better sex-targeted preventive and therapeutic strategies in SCC and other malignancies.

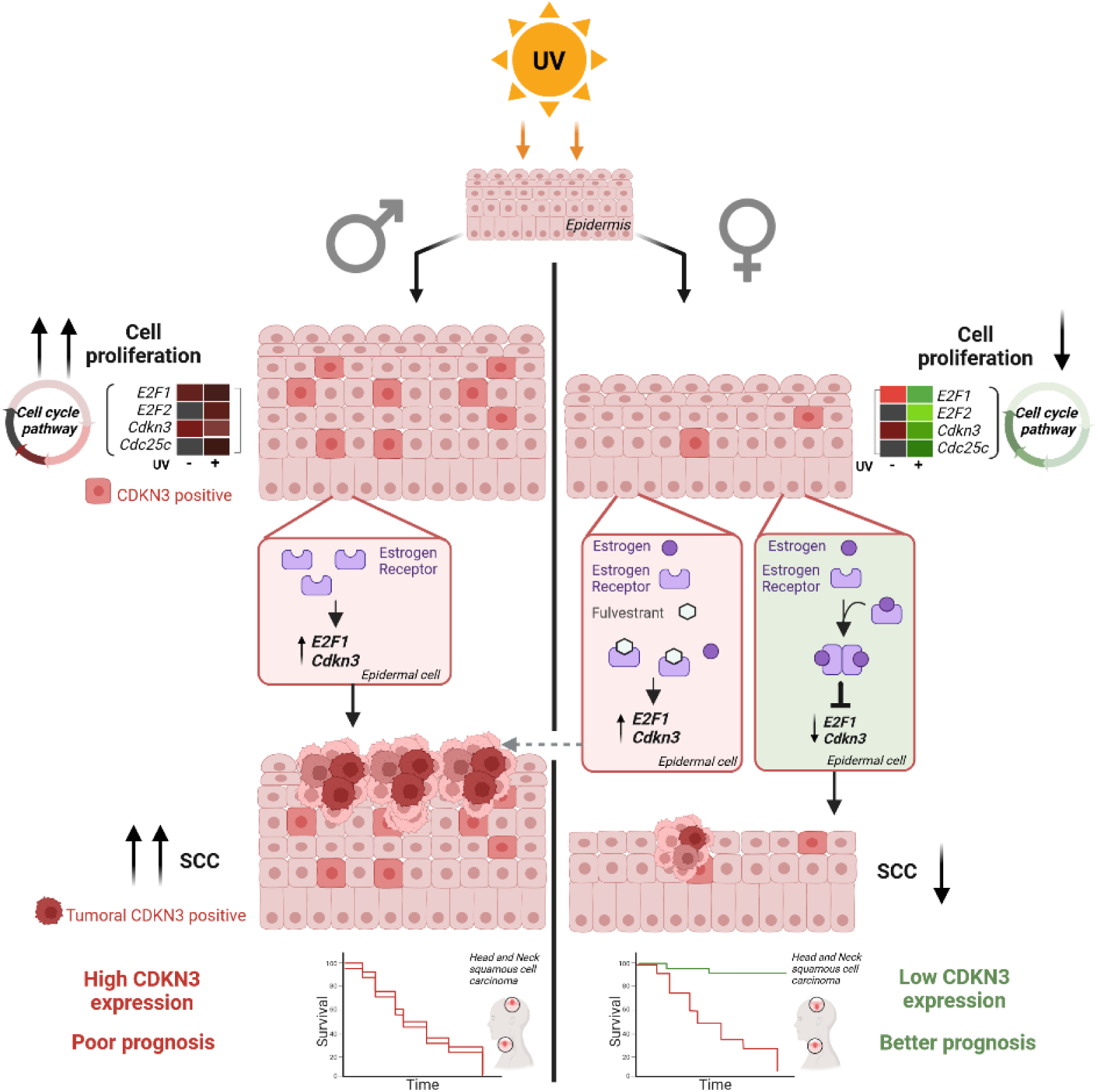

**Sex differences in UV-Induced cutaneous Squamous Cell Carcinoma: key insights into female cancer protection**

## Introduction

Differences in the incidence of at least 21 different cancers, excluding reproductive tissues, have been reported between men and women, with men having a greater risk of developing cancer than women^1,2^. This sex-difference also includes cancer mortality worldwide, with death rates 43% higher in men than in women^3^.

Cutaneous Squamous Cell Carcinoma (SCC), the second most common keratinocyte skin cancer^4^, is also affected by this sexual dimorphism. Indeed, sex is recognized as a risk factor for SCC as important as fair skin or advanced age^5^, and a recent meta-analysis of cutaneous head and neck SCC reported that men are more commonly affected than women^6^. This sex-disparity in SCC prevalence has been mainly attributed to differences in lifestyle (e.g. outdoor work^7^), behavior (e.g. motivation to sun protection measures and men shorter hair preferences^8^) and genetics (e.g. common men baldness^9^) related to UV exposure. Beyond sex-differences in the immune system^10–12^ and in oxidative stress in response to UV exposure^13^, intrinsic factors and physiological differences remain largely underexplored.

In particular, the role of sex hormones in modulating vulnerability to SCC is barely understood. Nevertheless, an elevated risk of developing keratinocyte skin cancers was associated with the use of oral contraceptives and with hormone replacement therapy^14,15^. Along the same lines, the androgen receptor seems to be a key determinant of melanoma development, another type of skin cancer^16^. Besides multiple genes present on sex chromosomes have been proposed to also play a role in oncogenesis^1^. Collectively, these data strongly suggest that sex hormones have a role in modulating SCC incidence in humans.

UV is the main carcinogenic factor for SCC^17,18^, but sex cellular and molecular responses of the skin to UV rays have not been described. UV provokes direct DNA damage which leads to mutations if unrepaired. UV exposure triggers a cascade of molecular events called the DNA damage response in exposed cells, aiming to repair DNA damage. Cell cycle arrest is required to allow a proper and timely regulated DNA repair and can switch to apoptosis if the DNA damage burden exceeds the repair capacity of the cell^19^. This initial response is followed by a mitogenic response, which leads to epidermal hyperplasia, a thickening which improves the protection of the basal layer of the epidermis against further penetration of UV rays^20,21^. The cutaneous response to UV also includes the production of various inflammatory cytokines such as Interleukin 1 (IL-1), inflammatory metabolites, arachidonic acid and mast cell-derived mediators leading to sun burn reaction^22,23^.

Understanding the role of sex hormones and their receptors, as well as sex-disparities in UV responses, is crucial to understand why women are better protected than men from the risk of developing SCC. Here, we provide cellular and molecular evidence underlying differences in SCC development in males and females using a relevant mouse model for UV-induced skin cancers. Specifically, we demonstrate that UV exposure triggers a sex-dependent response through the modulation of the E2F1/CDKN3 axis mediated by estrogen receptors. Finally, we prove the relevance of our observations to humans, using *ex vivo* skin explant cultures from women and squamous cell carcinoma biopsies from patients.

## Results

### Chronic UV exposure enhances squamous cell carcinoma development in males compared to females

Chronic UV exposure is a major risk factor for skin carcinogenesis. Men are more likely to develop skin cancer and the reasons for that are still poorly understood. We used a mouse model to better document such sex-disparities, comparing the skin responses of male and female mice to chronic UV exposure. B6.129-SKH1-Hr*^hr^* mice were exposed to UV three times per week for 6 months (UV 70mJ/cm^2^: a sub-erythematous dose), which provoked the formation of various lesions on their dorsal skin. Macroscopic observations revealed that the dorsal skin of males was far more affected than females: on average, 65% of the dorsal skin surface of males showed UV-induced lesions, compared to only 27% of the dorsal skin surface of females (Fig. 1a).

**Fig. 1.**
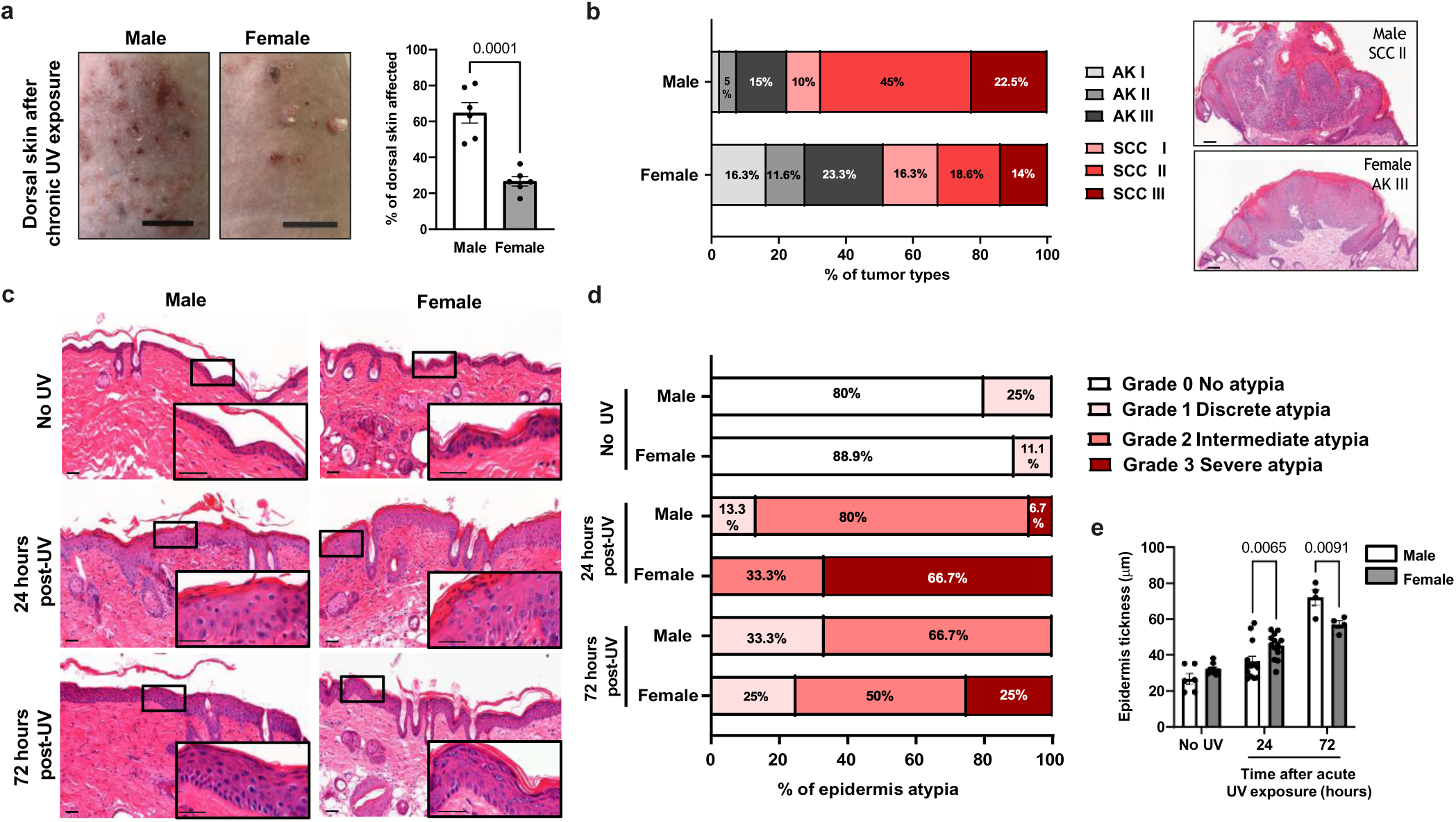
Chronic and acute UV exposure induce distinct, sex-specific skin responses in male and female mice. **a. Left:** Representative images of the dorsal skin in male and female mice following chronic UV exposure (3 times per week for 6 months; scale bar: 1 cm). **Right**: Percentage of dorsal skin with UV-induced lesions in male and female following chronic UV exposure. *n = 6 mice*, mean ± SEM, unpaired *t*-test. **b. Left:** Percentage of actinic keratosis (AK) and squamous cell carcinoma (SCC) lesions graded, based on lesions collected from male (n=43) and female (n=40) mice after chronic UV exposure. **Right:** Representative H&E-stained micrographs of the predominant tumor types for each group. Scale bar: 100 μm. **c.** Representative H&E-stained images of dorsal skin in male and female mice 24 hours or 72 hours after a single dose of UV exposure (120mJ/cm²), compared to control mice not exposed to UV (No UV). Scale bar: 50 μm and for the magnified image, 10 μm. **d.** Histogram showing the percentage of epidermal atypia grading in male and female mice following a single dose of acute UV exposure (120 mJ/cm²) at 24 hours and 72 hours time points, compared to control mice (No UV). *n= 4 to 12 mice*. **e.** Epidermal thickness measurements in male and female mice following a single dose of acute UV exposure (120mJ/cm²) at 24 hours and 72 hours time-points, compared to control mice (No UV). *n= 4 to 12 mice*, mean ± SEM, two-way ANOVA followed by Tukey’s multiple comparisons test.

The severity of these lesions was further characterized by a dermato-pathologist in a blinded manner. Chronic UV exposure of the skin is well-known to lead to the development of precursor tumoral lesions named actinic keratosis (AK) and of cancers of epithelial origin named cutaneous squamous cell carcinoma (SCC). Female mice presented, on average, higher number of benign AK lesions, but fewer malignant SCC lesions compared to male mice (Fig. S1a). Each collected AK and SCC was then further characterized according to its tumor grade, based on the degree of keratinization and of keratinocyte differentiation. This characterization revealed that tumors were more severe in males compared to females (Fig. 1b). Indeed, the commonest tumors collected in males were SCC grade II, while in females the commonest of the collected tumors were AK grade III (Fig. 1b). Regarding the most severe tumors, 22.5% versus 14% of the collected lesions were SCC III in males and females, respectively. As AK (grade III) and SCC (grade II) are the predominant tumor types in females and males respectively, we next investigated their proliferation rates by Ki67 staining. No difference between sexes was observed (Fig. S1b and c), and tumor depth increased with the severity of SCC grade, similarly in both sexes (Fig. S1d). In summary, male mice bear a greater overall tumor burden compared to females, with a predominance of more advanced lesions upon chronic exposure. This mouse model of photo-carcinogenesis therefore recapitulates epidemiological data in humans that cannot be attributed to lifestyle or behavior, making it an excellent model for studying the cellular and molecular basis of sex-disparity in the development of UV-induced SCC.

While chronic exposure to UV rays causes AK and SCC, acute UV exposure (or single UV exposure, 120mJ/cm^2^) induces transient damages resulting in reversible cellular atypia, which we then characterized in the epidermis of male and female mice. Atypia was graded by a dermato-pathologist in a blinded manner 24 hours and 72 hours after UV exposure, based on disorganization of epidermal layers and apparent phenotypical changes in histological sections (Fig. 1c). All epidermis samples showed some degree of atypia 24 hours after UV exposure, which diminished in severity after 72 hours, demonstrating skin recovery over time. Atypia severity was strikingly different between the sexes, affecting more females than males. Specifically, 66.7% of females versus only 6.7% of males presented a severe atypia (grade 3) 24 hours after UV exposure. At the same time point, most (80%) of males showed intermediate atypia, whereas 33.3% of females showed intermediate atypia. Differences between sexes remained 72 hours after UV exposure, with 25% of females still presenting severe atypia, a condition no longer present in males (Fig. 1d). Epidermal thickness increased in both sexes in response to acute UV (Fig. 1e). However, 72 hours after UV exposure, males presented greater epidermal thickening than females, with an average thickness of 72 μm for males versus 57 μm for females (Fig. 1e).

Collectively, these data show that both acute and chronic UV exposure affect the epidermis in a sex-specific way. On the long run, male mice develop more severe skin lesions caused by chronic UV exposure whereas, in the short term, female mice are more affected by a single dose of UV exposure.

### UV exposure induces similar DNA damages and mutations patterns in male and female epidermis

We next addressed the mechanisms underlying such sex disparities in the skin responses to UV. We first assessed if these differences arise from variations in direct DNA damage provoked by UV in male and female epidermal cells. Cyclobutane pyrimidine dimers (CPD) are the major type of direct DNA damage caused by UV exposure, representing 75 to 90% of the total UV lesions^24^. We observed similar amounts of CPD one hour after a single UV exposure (120mJ/cm^2^) in male and female epidermis, which were being efficiently removed over time by the DNA repair process underway in both sexes (Fig. 2a). Among the numerous proteins involved in the DNA damage response, the histone variant H2AX plays an important role. Upon DNA damage, H2AX is phosphorylated at Ser-139^25^ (γH2AX), which is the first step for localizing DNA damage and recruiting repair proteins. Using immunofluorescent staining of γH2AX as a biomarker for DNA damage, we observed that 24 hours after acute UV exposure, male and female murine epidermis exhibited a similar proportion γH2AX-positive keratinocytes presenting a foci staining pattern (Fig. 2b).

**Fig. 2.**
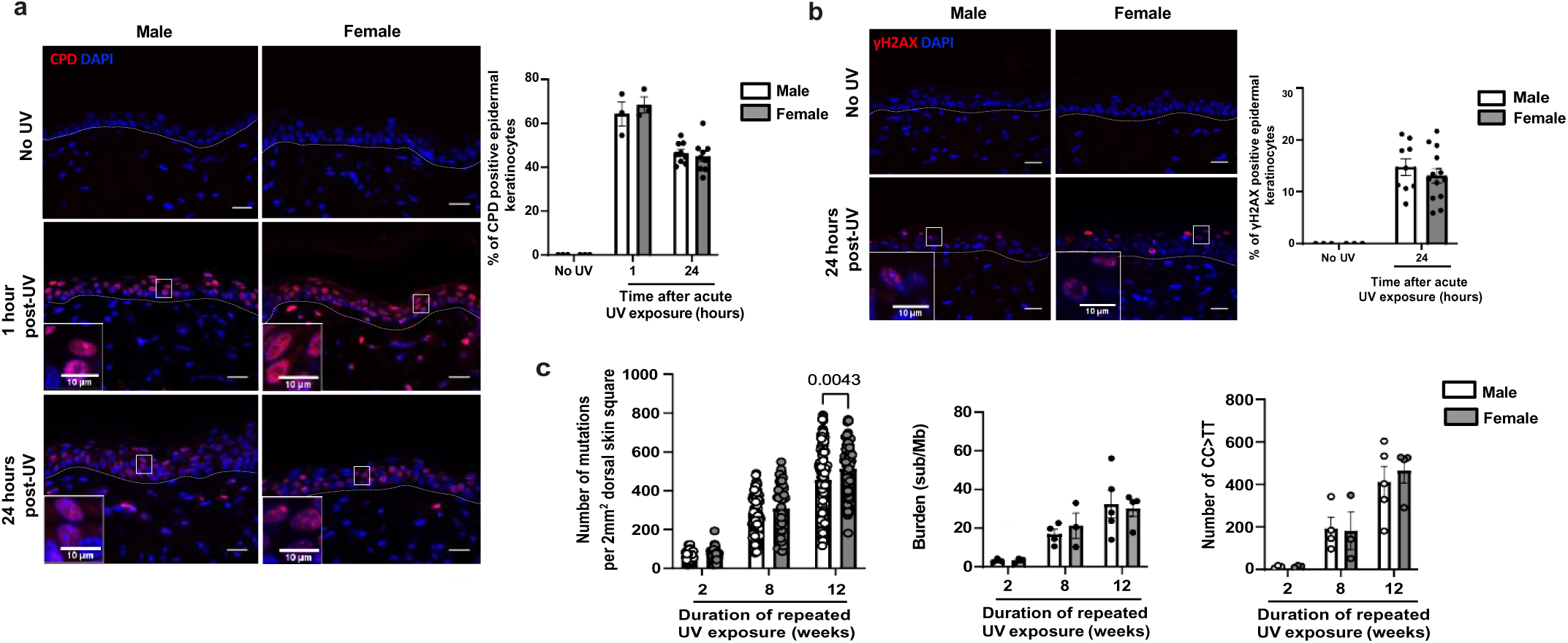
UV exposure induces similar level of epidermal DNA damage and mutations in males and female mice epidermis. **a. Left:** CPD (red) immunofluorescence in male and female dorsal epidermis collected 1 hour and 24 hours after a single dose of acute UV exposure (120mJ/c-m²), compared to control skin (No UV). DAPI was used as counterstaining (blue). The dotted line separates the epidermis from the dermis. Scale bar: 20 μm. **Right:** Percentage of CPD positive epidermal cells. *n(fields)= 4 per mouse, n= 3 to 9 mice*, mean ± SEM, two-way ANOVA. **b. Left:** γH2AX (red) immunofluorescence in male and female dorsal epidermis collected 24 hours after a single dose of acute UV exposure (120mJ/cm²), compared to control skin (No UV). DAPI was used as counterstaining (blue). The dotted line separates the epidermis from the dermis. Scale bar: 20 μm. **Right:** Percentage of γH2AX foci staining pattern positive epidermal cells. *n(fields)= 4 per mouse, n= 3 to 9 mice*, mean ± SEM, two-way ANOVA. **c. Left:** Number of mutations detected per 2 mm² in male and female dorsal epidermis exposed to UV (70mJ/cm²) for 2, 8, or 12 weeks, *n(square)= 19-22 per mouse*, *n= 3 to 5 mice,* mean ± SEM, two-way ANOVA with Sidak’s post hoc test. **Middle:** Mutation burden in male and female dorsal epidermis collected from mice exposed to UV (70mJ/cm²) for 2, 8, or 12 weeks, *n= 3 to 5 mice*, mean ± SEM, two-way ANOVA with Sidak’s post hoc test. **Right:** Total counts of CC>TT mutations in male and female dorsal epidermis collected 2, 8, or 12 weeks after multiple dose of UV exposure (70mJ/cm²), *n= 3 to 5 mice*, mean ± SEM, two-way ANOVA with Sidak’s post hoc test.

Mutations are acquired in response to external agents such as UV ^17^. We quantified the number of mutations present in 2mm^2^ areas of the epidermis of male and female mice exposed to UV for 2, 8 or 12 weeks followed by ageing, by ultradeep (average 1361x target coverage) targeted sequencing of 74 genes commonly mutated in SCC and other cancers^26,27^ (Fig. 2c). As expected, we observed that the total number of mutations per 2mm^2^ of epidermis increased with the duration of UV exposure, with a 7-fold increase between the groups exposed for 2 compared to 12 weeks (Fig. 2c, left). Along the same lines, the epidermal mutation burden, quantified as the number of synonymous substitutions per megabase, increased 3-fold between the groups exposed for 2 versus 12 weeks (Fig. 2c, middle). However, we observed no difference either in the total number of mutations or the mutation burden between male and female epidermis at any time point except for the total number of mutations 12 weeks after repeated UV exposure (Fig. 2c). We compared the mutational spectrum of our data to the mutational signatures SBS7a and SBS7b associated with exposure to UV light (COSMIC, Mutational Signatures). These signatures are characterized by cytosine substitution by thymidine (C>T) at TCN (SBS7a) or CCN/ TCN (SBS7b) trinucleotide sites, that may reflect different pyrimidine-dimer photoproducts induced by UV exposure (Fig. S2a). In analyzing these signatures, we observed no difference in the proportion of mutations assigned to UV associated signatures between the sexes (Fig. S2b). To collect more detailed information, we next quantified the CC to TT double base substitutions which are strongly associated with exposure to UV light. The number of CC to TT mutations increased with the duration of UV exposure at a similar rate between males and females (Fig. 2c, right). We also examined selection in this mutated landscape of synonymous, missense, nonsense and indel mutations, using dN/dS. Mutations in genes that confer a fitness advantage have a dN/dS ratio greater than 1 while those that are detrimental to cell fitness have a dN/dS ratio of less than 1. Examining the entire data set showed that several mutant genes were under positive selection (Fig. S2c), however comparing the strength of selection between sexes at the different time points showed no significant differences (Fig. S2d).

Overall, these data show that the differences in atypia, tumor development and tumor severity observed in males and females after UV exposure cannot be explained by differences in UV-induced DNA lesions or mutation patterns, nor by the nature of the mutant genes that are selected.

### UV exposure affects epidermal proliferation in males, while it affects differentiation in females

It has been shown that exposure of the skin to UV rays affects the rate of epidermal keratinocyte proliferation and the proportion of dividing cells^20^, a response which must always be balanced with differentiation to maintain epidermal homeostasis^29^. We then asked if an imbalance between proliferation and differentiation could be involved in the contrasting UV response in male and female epidermis described in Fig. 1. After a single acute UV exposure (UV 120mJ/cm2), we quantified epidermal proliferation in UV-exposed male and female epidermis, using Ki67 labeling of skin sections (Fig. 3a). Examination of control samples showed that background cell proliferation was similar in male and female epidermis, with 24.8% and 27.4% of Ki67-positive epidermal cells, respectively. We observed a proliferative response of epidermal layers in two steps. First, an early response 24 hours after UV exposure, when we quantified a significant decrease in epidermal cell proliferation, which was similar in male and female epidermis (Fig. S3a). Second, a later response 72 hours after UV exposure, when we observed a strong increase in epidermal cell proliferation in male epidermis only, while in females, the proliferation rate returned to the unexposed control (Fig. 3a). Of note, while the proliferation rate had returned to control level 72 hours after UV exposure in female epidermis, males did not turn off proliferation in epidermal suprabasal layers (spinous layer specifically) at that time point (Fig. 3a). This proliferation of keratinocytes in the layers above the basal layer is a characteristic feature of AK^30^. At the mRNA level, these results were confirmed by an increase of *Mki67* mRNA expression only in male epidermis (Fig. S3b).

**Fig. 3.**
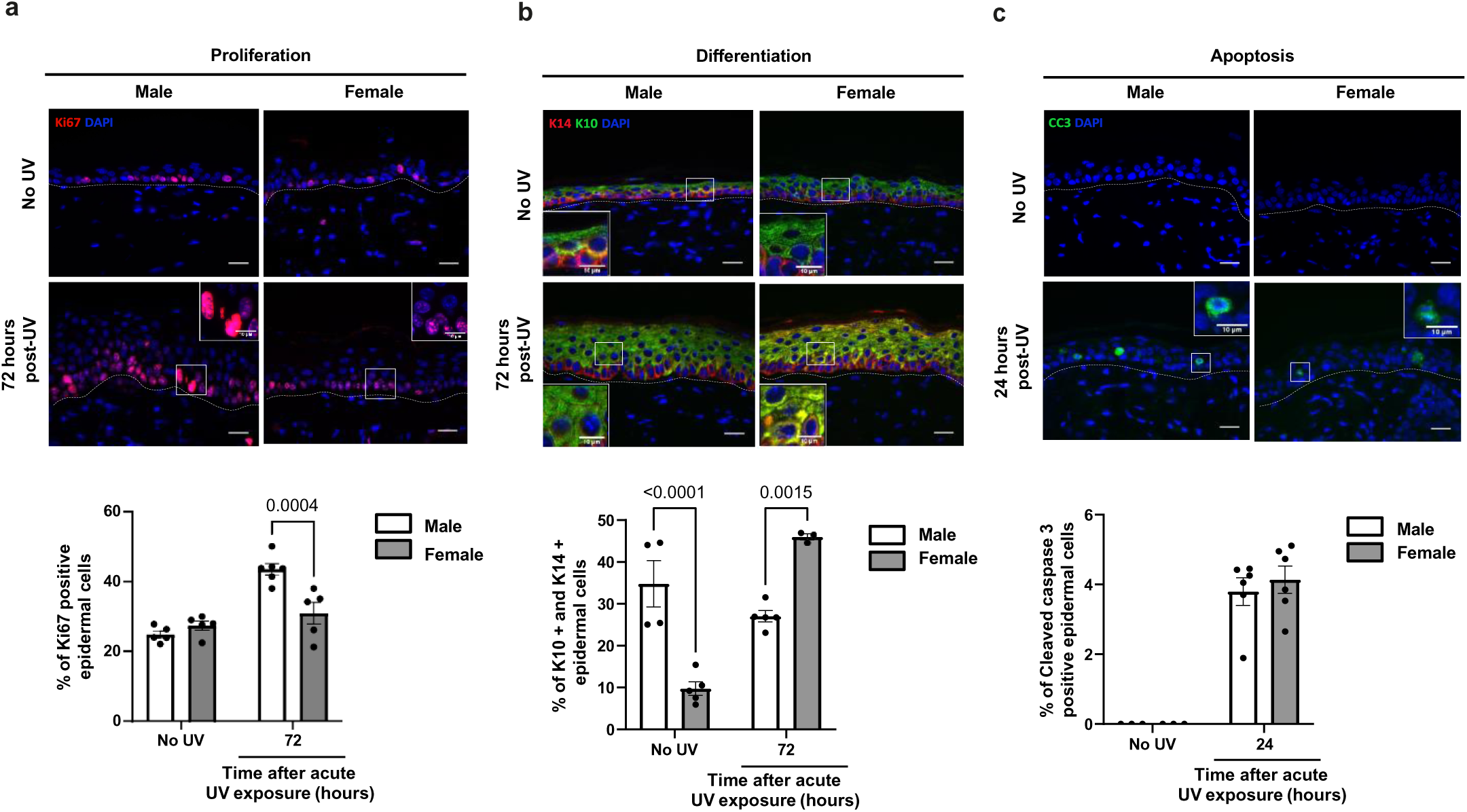
UV exposure affects epidermal proliferation in males and differentiation in females mice. **a. Top:** Ki67 (red) immunofluorescence in male and female dorsal epidermis collected 72 hours after a single dose of acute UV exposure (120mJ/cm²), compared to control skin (No UV). DAPI was used as counterstaining (blue). The dotted line separates the epidermis from the dermis. Scale bar: 20 μm. **Bottom:** Percentage of Ki67 positive keratinocytes. *n(fields)= 4 per mouse, n= 5 mice*, mean ± SEM, two-way ANOVA with Sidak’s post hoc test. **b. Top:** Keratin 14 (K14; red) and Keratin 10 (K10; green) immunofluorescences in male and female dorsal epidermis collected 72 hours after a single dose of acute UV exposure (120mJ/cm²), compared to control skin (No UV). DAPI was used as counterstaining (blue). The dotted line separates the epidermis from the dermis. Scale bar: 20 μm. **Bottom:** Percentage of double K10-positive and K14-positive epidermal keratinocytes. *n(fields)=4 per mouse, n= 3 to 6 mice*, mean ± SEM, two-way ANOVA with Sidak’s post hoc test. **c. Top:** Representative micrographs of cleaved caspase3 (CC3; green) immunofluorescence in male and female dorsal epidermis collected 24 hours after a single dose of acute UV exposure (120mJ/cm²), compared to control skin (No UV). DAPI was used as counterstaining (blue). The dotted line separates the epidermis from the dermis. Scale bar: 20 μm. **Bottom:** Percentage of CC3-positive keratinocytes and quantification. *n(fields)= 4 per mouse, n= 3 to 6 mice*, mean ± SEM, two-way ANOVA with Sidak’s post hoc test.

In healthy skin, epidermal homeostasis is supported through a well-balanced equilibrium between proliferation and differentiation. Keratinocytes proliferate in the basal layer of the epidermis, then they exit cell cycle and undergo differentiation while migrating up into the suprabasal layers ^31,32^. We next studied the proliferation/differentiation equilibrium in response to UV exposure, using Keratin 14 (K14) and Keratin 10 (K10) as prototypic markers to label basal/dividing and suprabasal/non-dividing layers of keratinocytes, respectively (Fig. 3b). As suggested by both morphology and the percentage of cells expressing only K14 (K10 negative and K14 positive), neither UV exposure nor sex seems to affect the keratinocytes of basal layer (Fig. S3c-d). The co-expression of K14 and K10 is usually considered as abnormal and the expansion of the expression of the basal keratinocyte marker K14 to suprabasal layers suggests a failure of keratinocytes to shut down K14 expression upon differentiation. In absence of UV, we saw that male epidermis presented a greater proportion of keratinocytes expressing both K14 and K10 compared to female epidermis (Fig.3b). In response to UV, we observed an increase of K10 and K14 marker co-expression in the suprabasal layers both in male and female epidermis (Fig. S3c). Importantly, the abnormal co-expression of K10 and K14 persisted and increased 72 hours after UV exposure only in female epidermis (Fig. 3b), which suggests that the return to a normal differentiation process differs in female epidermis. In contrast, increased co-expression of K14 and K10 in response to UV was transient in male epidermis (Fig. S3c). In agreement with this observation, expression of the K10 marker was decreased after UV exposure only in females (Fig. S3d).

UV exposure is also well-known to provoke apoptosis ^33^, which is also involved in the maintenance of epidermal homeostasis. To quantify apoptosis in UV-exposed epidermis, we performed Cleaved caspase 3 (CC3) staining on histological sections of male and female epidermal skin 24h after a single UV exposure. We observed no difference in the number or localization of apoptotic epidermal cells (CC3-positive) between male and female epidermis (Fig. 3c).

In brief, we discovered that two major cellular responses to UV exposure are differentially influenced by sex. In males, UV exposure primarily impacts proliferation, whereas in females, it predominantly affects epidermal differentiation.

### Sex-specific modulation of epidermal cell cycle and E2F transcription factors in response to UV

Our analyses so far show intrinsic differences in the epidermal response to acute and chronic UV exposure between male and female mice in terms of tumor incidence and progression, as well as proliferation and differentiation. To investigate the molecular mechanisms involved in this sex-specific response to UV, we analyzed the transcriptome of male and female mouse epidermis exposed to UV using RNA sequencing (four epidermal samples per sex, non-UV exposed controls and 24 hours after UV exposure) (Fig 4a). The 24 hours post UV exposure time point was chosen based on *i)* the peak of atypia in both males and females (Fig. 1) and *ii)* an increase of *Il1b* and *Tgfb-1* expression, two cytokines known to be induced in response to UV exposure and used here as a marker of effective epidermal response^34^ (Fig. S4). Using hierarchical clustering, differentially expressed genes in male (Fig. S5) and female (Fig. S6) epidermis were mapped. 9658 and 9703 genes were identified as differentially regulated in response to a single dose of UV in male and female epidermis, respectively (adjusted p-value< 0.05). Differentially regulated genes were grouped into three clusters in male epidermis, with clusters I and II including down-regulated genes and cluster III up-regulated genes in response to UV (Fig. S5). Four clusters of differentially regulated genes were identified in female epidermis: clusters I and II with down-regulated genes and clusters III and IV with up-regulated genes in response to UV (Fig. S6). Reactome Pathway Enrichment analysis of differentially expressed genes revealed that several pathways were differentially regulated by UV exposure in male and female epidermis. Most of the genes with UV-induced upregulated expression were similar in male and female epidermis and were enriched in “ribonucleoprotein complex biogenesis”, “mitochondrial gene expression”, “regulation of apoptotic signaling” and “keratinocyte differentiation” pathways (Fig. S5 and S6). The only exception was the “response to oxidative stress” pathway that was up-regulated only in male epidermis in response to UV. In contrast, the genes whose expression was reduced by UV exposure were mainly sex-specific. “Mitotic cell cycle phase transition”, “positive regulation of cell cycle”, “double strand break repair” and “cell cycle checkpoint signaling” were the most significantly down-regulated pathways only in the epidermis of female mice (Fig. S6). In male epidermis, the two most down-regulated pathways were “lymphocyte differentiation” and “regulation of T cell activation” (Fig. S5).

**Fig. 4.**
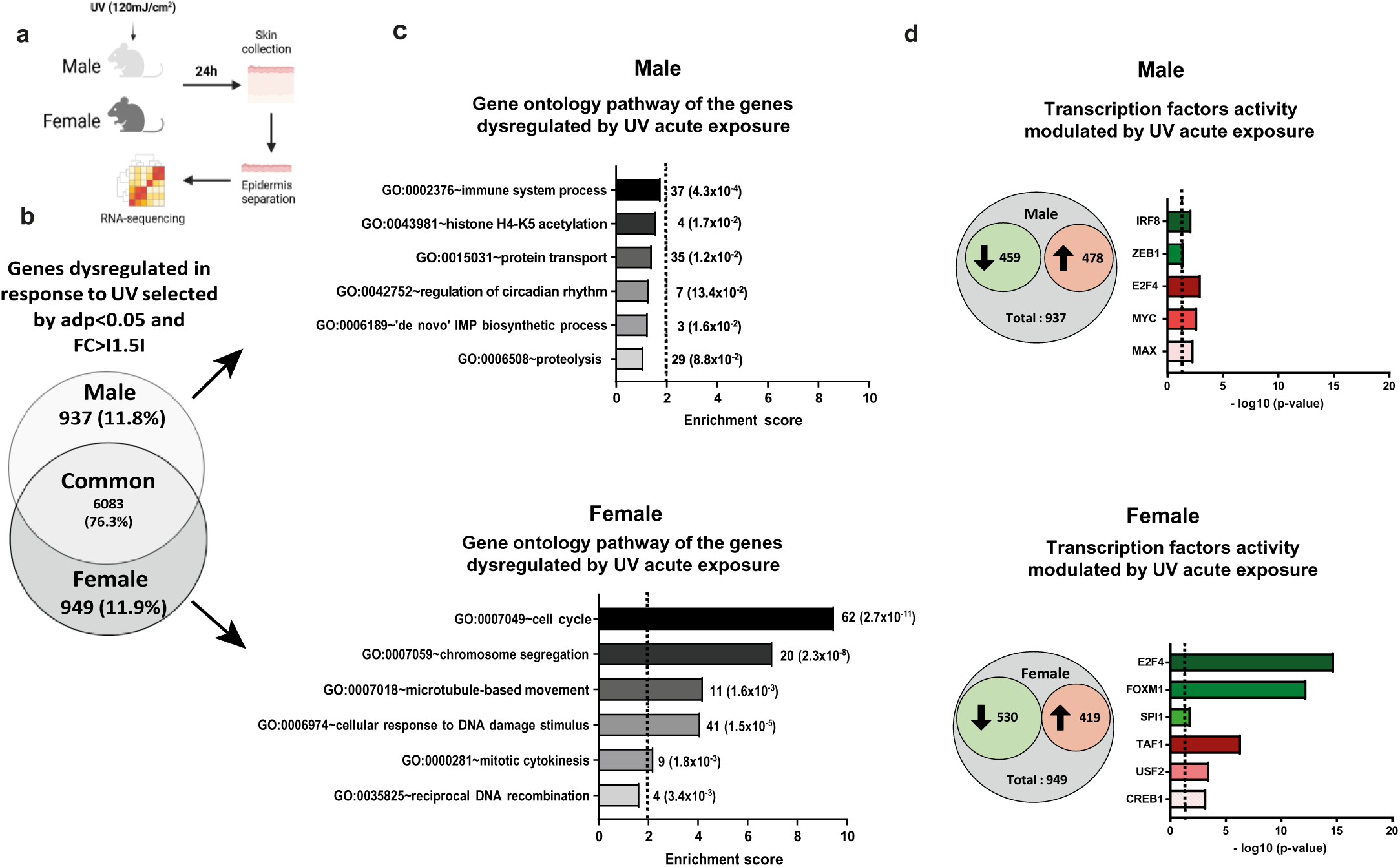
Acute UV exposure triggers female-specific downregulation of cell cycle pathway and E2F in the epidermis. **a.** Experimental flowchart. **b.** Venn diagram illustrating the number of genes with significant changes in expression (adjusted p-value <0.05 and FC>I1.5I) in male and female mouse epidermis 24 hours post-UV exposure (single dose; 120mJ/cm²). **c.** Gene ontology pathway analysis of the genes significantly dysregulated in response to acute UV exposure specifically in male, 937 genes (top) and in male, 949 genes (bottom) samples. Histograms show the enrichment score for each identified pathway. Gene number and the corresponding *p*-value are indicated to the right oh the histograms. **d.** Histograms showing the number of downregulated genes - 459 in males and 530 in females - and upregulated genes - 478 in males and 419 in females - specifically in male (top) or female (bottom) samples, accompanied by transcription factor enrichment analysis of the dysregulated genes in each sex.

A Venn diagram of the differentially expressed genes (DEGs; adjusted p-value< 0.05; Fold change IFCI>1.5), illustrates that the majority of DEGs were common for both male and female epidermis (6083, 76.3%) (Fig. 4b), while approximately 12% of the genes exhibited a sex-specific response to UV (937 and 949 in male and female epidermis, respectively). Gene ontology analysis of these sex-specific DEGs revealed that a majority belonged to “immune system process” (37), “protein transport” (35) and “proteolysis” (29) in male epidermis (Fig. 4c) and to “cell cycle” (62) and “cellular response to DNA damage stimulus” (41) in females epidermis (Fig. 4c). We then focused on the most significant groups of genes, with the highest enrichment score and showing a sex-specific regulation, that is the group of 62 genes dysregulated upon UV exposure in female epidermis only, belonging to “cell cycle biological process” (GO:0007049; enrichment score of 9.5). To identify transcription factors associated with this group of DEGs, we analyzed the up- and downregulated sex specific genes separately referring to ENCODE^35^ and ChEA^36^ Consensus TFs from ChIP-X category in Enrichr^37^ (Fig. 4d). In male epidermis exposed to UV, we found an enrichment of upregulated genes associated with three transcription factors namely E2F4, the proto-oncogene MYC and MAX, while downregulated genes were associated with the two transcription factors IRF8 and ZEB1 (Fig. 4d). In female epidermis exposed to UV, upregulated genes were associated with the three transcription factors TAF1, USF2 and CREB1, while downregulated genes were associated with the three transcription factors E2F4, FOXM1 and SPI1 (Fig. 4d). The most significant enrichment was observed for E2F4 where 60 out of its 710 target genes where among the downregulated genes in UV exposed female epidermis (*p*-value: 1.9×10^-16^).

Overall, we identified that 12% of the total genes deregulated in response to UV are sex-specific. Importantly, we identified the transcription factor E2F, a major regulator of the cell cycle whose activity is repressed in response to UV specifically in female epidermis.

### Estrogen receptors-mediate UV-induced modulation of the proto-oncogene CDKN3 in the epidermis of female mice and in squamous cell carcinoma

The E2F family of transcription factors particularly caught our attention as key regulators of the cell cycle, for which sex-specific regulation could explain the sex-specific proliferation and differentiation responses that we observed following UV exposure (Fig. 3). There are seven subgroups in the E2F family. E2F1-3 are required for normal cell cycle progression, while in contrast E2F4 and 5 are mainly involved in cell cycle exit and differentiation. The roles of E2F6 and 7 are still unclear, but they are believed to act as transcriptional repressors of the cell cycle^38^. In quantifying the expression of the transcription factors themselves, we found that *E2f1* and *E2f2* gene expression was decreased in response to UV only in female epidermis (Fig. 5a and Fig. S7a), while the expression of *E2f3*, *E2f4*, *E2f5*, *E2f6, E2f7* genes was not affected by sex (Fig. S7a). In line with these observations, the expression of the E2F target genes, the proto-oncogene *Cdkn3* and *Cdc25c*, which are mainly involved in cell cycle progression, were also down-regulated in response to UV only in female epidermis (Fig. 5a and Fig. S7b).

**Fig. 5.**
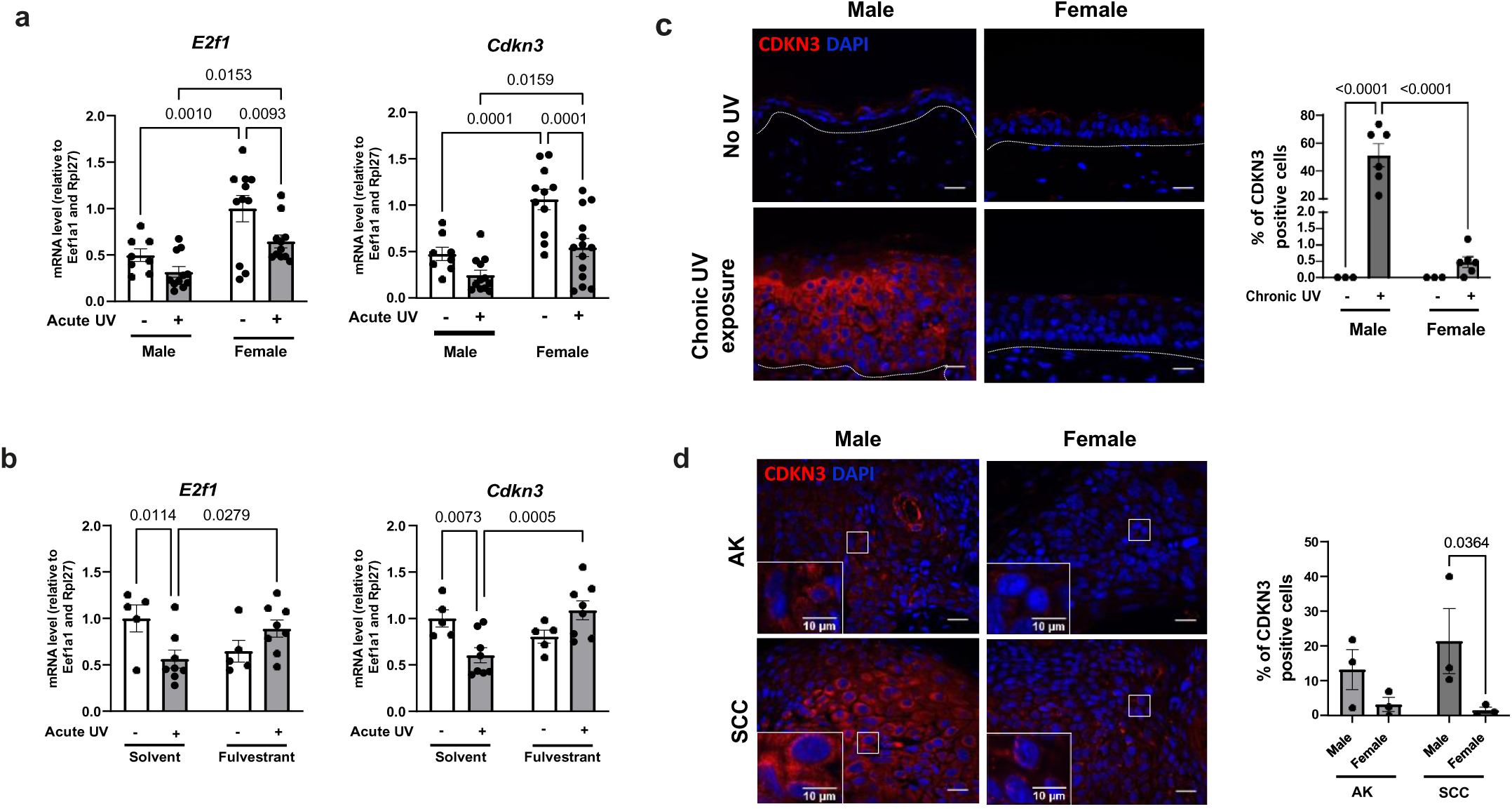
Estrogen receptor-mediated UV-induced downregulation of the proto-oncogene CDKN3 in the female epidermis and in squamous cell carcinoma. **a.** Quantification of the relative gene expression of *E2f1*, *Cdkn3* at mRNA level by RT-qPCR in epidermal samples from male and female mice. *n=8-14 mice per sex*, mean ± SEM, two-way ANOVA with Sidak’s post hoc test. **b.** Quantification of the relative gene expression of *E2f1* and *Cdkn3* at the mRNA level by RT-qPCR in epidermal samples from female mice treated with solvent or fulvestrant (150 mg/kg) for 48 hours prior to acute UV exposure. Data represent the mean values for *n=5 mice* (solvent) and *n=8 mice* (fulvestrant), mean ± SEM, two-way ANOVA with Sidak’s post hoc test. **c. Left:** CDKN3 (red) immunofluorescence in male and female dorsal epidermis following chronic UV exposure (3 times per week for 6 months). DAPI was used as counterstaining (blue). The dotted line separates the epidermis from the dermis. Scale bar: 20 μm. **Right:** Percentage of CDKN3 positive epidermal keratinocytes. *n(fields)=4 per mouse*, *n= 3 to 6 mice*, mean ± SEM, two-way ANOVA with Sidak’s post hoc test. **d. Left:** CDKN3 (red) immunofluorescence in actinic keratosis (AK) and squamous cell carcinoma (SCC) biopsies collected from male and female mice following chronic UV exposure. DAPI was used as counterstaining (blue). Scale bar: 20 μm. **Right:** Percentage of CDKN3 positive cells. *n(fields)=4 per mouse, n= 3 tumors per sex,* mean ± SEM, two-way ANOVA with Sidak’s post hoc test.

To test whether the downregulation of the female-specific genes *E2f1*, *E2f2*, *Cdkn3*, and *Cdc25c* in response to acute UV exposure was dependent on estrogen receptors, female mice were treated with fulvestrant, a pharmacological estrogen receptor antagonist, 24 hours prior to UV exposure. As expected, we observed a reduction in uterus weight in the fulvestrant-treated mice (Fig. S8a) and an increase in *Il1b* gene expression (Fig. S8b), indicating the effectiveness of estrogen receptor antagonism and UV exposure, respectively.

Consistent with our data, we found that UV exposure downregulated the expression of *E2f1, E2f2, Cdkn3*, and *Cdc25c* in the solvent control condition (Fig. 5b and Fig. S8c). Significantly, when estrogen receptors were degraded following fulvestrant treatment, this downregulation was abolished for *E2f1* and *Cdkn3*, the typical UV-induced decrease was entirely lost, demonstrating that the UV-induced reduction in the expression of these genes is mediated by estrogen receptors (Fig. 5b).

To assess whether the acute effects of UV exposure on CDKN3 persist after chronic exposure, we measured CDKN3 expression in chronically UV-exposed, non-pathological mouse skin in both males and females. We observed an increase in epidermal CDKN3 protein expression only in males in chronically irradiated skin compared to females (Fig. 5c). We further investigated these observations in pre-lesional actinic keratosis (AK) tissues and squamous cell carcinoma (SCC) tumors derived from male and female mice chronically exposed to UV. In SCC samples, we observed a significant increase in CDKN3-positive cells in males compared to females, with males exhibiting 15 times more CDKN3-positive cells than females (Fig. 5d). In AK biopsies, a slight increase in CDKN3 expression was detected in males compared to females (Fig. 5d).

As key findings, we identified that the expression of E2F1 and CDKN3 - key regulators of the cell cycle, with E2F1 functioning as a major transcription factor and CDKN3 as a proto-oncogene - was significantly reduced in response to UV exposure, but only in females. Mechanistically, this female-specific UV-induced downregulation of E2f1 and Cdkn3 is mediated by estrogen receptors. Notably, in SCC tissues, we observed a persistent sex-specific difference in CDKN3 protein expression, with significantly higher levels in males compared to females.

### Protective downregulation of the CDKN3 proto-oncogene in the skin of healthy women and in squamous cell carcinoma

The above findings suggest that differences in susceptibility of men and women to UV-induced skin malignancies is routed in biological and physiological differences, rather than behavioral differences. To go beyond our mouse model and to test this hypothesis in human skin, we performed a series of experiments on *ex vivo* human skin explants, and we analyzed malignant skin lesions from men and women patients. First, we assessed the responses of skin explants with phototypes I and II (Fitzpatrick scale) harvested from healthy women having abdominoplasty procedures, collected 24 hours and 72 hours after single increasing doses of UV (either 240, 480, or 720mJ/cm^2^). UV exposure induced a dose-dependent inflammatory response in skin explants as measured by the expression of *IL1b*, confirming effective epidermal response to UV (Fig. S9a). We then assessed the effect of UV exposure on proliferation using the immunostaining of nuclear Ki67. UV exposure led to a decrease in proliferation 24 hours and 72 hours after UV exposure (Fig. 6a), in line with the reduction in epidermal proliferation we also observed in female mouse epidermis (Fig. 3a). The expression of *E2F1*, *E2F2*, *CDKN3*, *CDC25C* mRNA was also reduced in a UV dose-dependent manner, with *E2F2* showing a 2.7-fold decrease in response to the highest dose of UV (Fig. 6b and Fig. S9b).

**Fig. 6.**
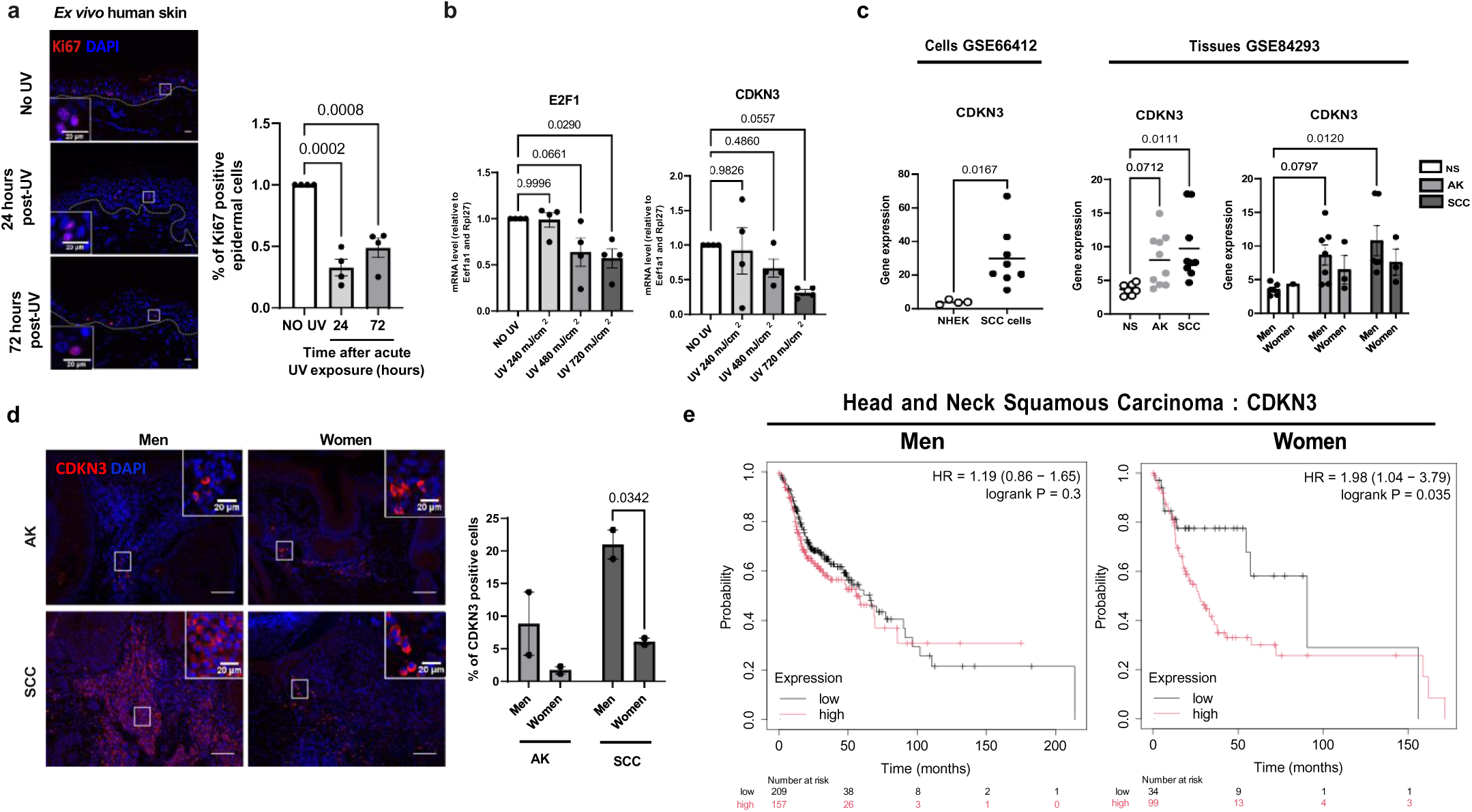
Downregulation of CDKN3 proto-oncogene expression in the skin of healthy women and in squamous cell carcinoma. **a. Left:** Ki67 (red) immunofluorescence in human *ex vivo* skin explant cultures from woman subjects, 24 hours and 72 hours after exposure to a single dose of UV (720mJ/cm²), compared to non-UV-exposed control explants. DAPI was used as counterstaining (blue). The dotted line separates the epidermis from the dermis. Scale bar: 50 μm. **Right:** Percentage of Ki67 positive epidermal cells. *n(field)=4 per subject, n= 4*, Mean ± SEM, two-way ANOVA with Tukey’s post hoc test. **b.** Relative mRNA expression of *E2F1* and *CDKN3* in *ex vivo* human skin explant cultures from woman subjects, 24 hours after exposure to UV with increasing intensities (240, 480 and 720 mJ/cm2), compared to non-UV-exposed control explants, quantified by RT-qPCR. *n=4 subjects*, Mean ± SEM, two-way ANOVA with Tukey’s post hoc test. **c. Left:** Expression of *CDKN3* mRNA expression in normal human epithelial keratinocytes (NHEK) and human Squamous Cell Carcinoma (SCC) from public datasets (GSE66412). *NHEK: n= 4, SCC: n= 8.* Mean ± SEM, t-student test. **Right:** Expression of *CDKN3* mRNA expression in normal human skin (NS), actinic keratosis (AK) and squamous cell carcinoma (SCC) lesions from men and women in public datasets (GSE84293). *NS: n= 7 (NS; 6 men and 1 woman), n= 10 (AK; 7 men and 3 women), n=9 (SCC; 5 men and 3 women)*. The left graph shows *CDKN3* expression with sex separation. Mean ± SEM, one-way ANOVA with Tukey’s post hoc test for non-sex-separated graph, two-way ANOVA with Tukey’s post hoc test for sex-separated graphs. **d. Left:** CDKN3 (red) mmunofluorescence in AK and SCC biopsies collected from men or women patients. DAPI was used as counterstaining (blue). Scale bar: 100 μm. **Right:** Percentage of CDKN3 positive cells. *n=2 tumors per sex*, for each 3 fields were counted. Mean ± SEM, two-way ANOVA with Sidak’s post hoc test. **e.** Kaplan-Meier survival curves for men (left) or women (right) patients with Head and Neck Squamous Cell Carcinoma (HNSC), based on *CDKN3* gene expression by Kaplan Meier plotter.

We then proceeded with a retrospective analysis of the genomic data in two independent sets of human samples. The GSE66412^39,40^ set of data includes 8 cutaneous SCC cell lines and 5 normal human epidermal keratinocytes (NHEK) cell lines; the GSE84293^41^ patient cohort includes Normal Skin (NS), Actinic Keratosis (AK) and Squamous cell carcinoma (SCC). In line with the experimental data we collected, SCC cell lines exhibited overexpression of *CDKN3* and *CDC25C* mRNA compared to healthy NHEK cells (Fig. 6c and Fig. S9c). This finding was confirmed in the second dataset, in which we observed a significant increase in the expression of *CDKN3* and *CDC25C* mRNA in SCC compared to healthy skin. While there was an increasing trend in the expression of *CDKN3* and *CDC25C* mRNA in AK compared to healthy skin, too, this was however not significant. Of much interest, the increased expression of *CDKN3* and *CDC25C*, was more prominent in men SCC lesions compared to women (Fig. 6c and Fig. S9d).

We next deepened these observations by analyzing the expression of CDKN3 protein in additional AK and SCC lesions derived from men and women patients. CDKN3 staining of tissue sections of men and women AK (*n=2*) and SCC (*n=2*) biopsies revealed that men lesions had 3-times more CDKN3-positive cells than women lesions (Fig. 6d). Moreover, the increase in CDKN3-positive cells in SCC compared to AK was more prominent in men that in women tissues sections.

Finally, we investigated whether the expression level of the E2F target genes *CDKN3* and *CDC25C* was correlated with patient survival rates. To do so, because such data are not available for cutaneous SCC, we turned to another form of carcinoma, highly aggressive, head and neck squamous cell carcinomas (HNSC). Available public databases, show that the expression of *CDKN3* and *CDC25C* genes in patients with HNSC, is negatively correlated with patient survival rate only in women, and not in men (Fig. 6e and Fig. S9e)^42^.

Overall, our data suggest that CDKN3 and CDC25C play a role in the difference in susceptibility between men and women to UV-induced skin cancers. Moreover, their lower expression in women, might have a protective effect against the development of skin SCC.

## Discussion

Our study provides new insights into intrinsic differences between male and female epidermal responses to both acute and chronic UV exposure, at the tissue, cellular and molecular levels. Our data emphasize that extrinsic factors, such as behavior ^8^, are not the sole contributors to the sex-disparity in SCC incidence. Indeed, chronic UV exposure leads to the development of more numerous and aggressive tumors in male compared to female mice, in line with findings from epidemiological studies in human^5^.

Epidermal proliferation is the most parameter we found differentially regulated with UV between male and female murine skins. Indeed, proliferation is greater in males in response to a single dose of UV, consistent with previous similar observations made with another carcinogenic factor (DMBA/TPA; less relevant for human cancers, however)^10^. Prolonged epidermal proliferation and hyperplasia are sufficient to promote the formation of early premalignant actinic keratosis (AK) lesions^43^. Along the same lines, UV activates epidermal proliferation and hyperplasia through activation of the epidermal growth factor receptor (EGFR)^20^. EGFR inhibition thus efficiently suppressed UV-induced epidermal proliferation, ultimately decreasing tumorigenesis ^44^. These data illustrate that the regulation of epidermal proliferation in response to UV is a critical factor of UV-induced cancer development. In the long term, this sex-dependent proliferative response to UV is likely crucial for sex-specific SCC development.

At the molecular level, we further discover that this sex-specific difference in epidermal proliferation is underpinned by an estrogen receptor-dependent and sex-specific regulation of the activity of the E2F1 transcription factor. The crucial role of E2F1 in controlling keratinocyte growth is well-illustrated by data showing that E2F1^-/-^ mouse primary keratinocytes cannot be maintained in culture^45,46^. In contrast, high expression and activity of E2F transcription factors were observed in many tumors and are usually associated with oncogenesis and poor prognosis^47–49^. In the skin, E2F was identified as a key transcriptional driver of squamous cell carcinoma development, whose activity is increased during the transition from healthy skin to actinic keratosis^41^. Meanwhile, in skin cutaneous melanoma, low levels of E2F1 and E2F2 were associated with better prognosis^50^. E2F1 may be thus considered as a potential target for the treatment of squamous cell carcinoma^51^. In support of the female-specific regulation of E2F1, existing research indicates that estrogen receptor alpha promotes E2F1 expression^52,53^ and estradiol regulates the expression of several E2F family members in breast cancer cell line MCF7^54^. Our work reveals that E2F1 exhibits decreased expression in response to UV only in the epidermis of female mice, identifying this transcription factor as a major player in the sex-specific difference of SCC development. Moreover, we discover that lower expression and activity of E2F are associated with the development of less numerous and less aggressive SCCs in female mouse epidermis, suggesting a protective role of low E2F activity against SCC development in females.

Among the target genes of E2F transcription factors, we identify the proto-oncogene CDKN3 as a marker exhibiting sex-specific regulation in response to UV in both healthy epidermis and SCC. CDKN3 is a cell cycle regulator, frequently overexpressed in various cancers and recognized as a tumor promoter^55–57^. Its high expression is associated with poor overall survival across eight types of cancers^58^. In skin cancers, CDKN3 has been identified as a marker of progenitor cells, linked to the suppression of epidermal differentiation and concurrent progression of SCC. Furthermore, its overexpression was associated with poorly differentiated SCC and worse prognosis^59^. Here, we show that UV-induced downregulation of the proto-oncogene CDKN3 occurs specifically in the female epidermis of both mice and humans. Importantly, this downregulation is mechanistically mediated by the estrogen receptor. This finding suggests a potential mechanism whereby the balance between CDKN3-expressing progenitor cells and epidermal differentiation is regulated in a sex-specific manner. This regulation likely plays a critical role in the sex-specific progression and outcomes of SCC. These insights highlight the importance of considering sex as a biological variable in understanding the molecular underpinnings of skin cancer progression.

In conclusion, we identified the epidermal markers E2F1 and CDKN3 as key players in the sex-specific protection observed in females against SCC development, both in mice and humans. These findings shed light on the molecular basis of sex dimorphism in skin cancer susceptibility.

## Methods

### Animal experiments

All experiments involving animals were approved by the Veterinary Office of the Canton Vaud (Switzerland) in accordance with the Federal Swiss Veterinary Office Guidelines and conform to the Commission Directive 2010/63/EU.

SKH-1 hairless mice (Crl: SKH1-Hr*^hr^*, Charles River) were crossed with Sv129/C57BL6/J male and female mice to obtain B6.129-SKH1-Hr*^hr^* mouse. This model is well established in the laboratory as part of our research into the role of PPARβ in the development of skin cancers and is therefore used for all projects^60^. Mice were raised, housed, and experienced in the conventional animal facility of the Center for Integrative Genomics, at the University of Lausanne. They were kept on IVC cages in a standard colony (2-5 animals per cage), in a light-controlled environment (12/12-h light/dark cycle, artificial light with daylight spectrum at an average intensity of 100 lux), with a hygrometry between 45% and 65% and a temperature of 22°C (+/-2 °C). They were housed on Aspen bedding (Safe select) and fed ad libitum with Sp-150 irradiated pellets (Safe) and filtered water.

For acute and chronic UV exposure, mice were UV irradiated on their backs with a GL40E 40W tube (SNEE), which emits most of its energy within the UVB range (90%; emission spectrum 280– 370; 10% UVA). Doses of UVB (312 nm) and UVA (370 nm) were monitored using an appropriate radiometer. The exposures were carried out in a (home-made) box where the mice were separated individually to avoid bias due to grouping, and the experiment took place in the animal facility.

Acute UV exposure: Females and males B6.129-SKH1-Hr*^hr^* aged 10-14 weeks old were placed in individual room and irradiated on their backs with a UV lamp. UV radiation was monitored using a radiometer until a dose of 120mJ/cm2 was reached in approximately 3min. This is a sub-erythematous dose. Control mice were sham manipulated but not exposed to UV. Mice were euthanized 1 hour, 24 hours or 72 hours after UV exposure by lethal intraperitoneal injection of pentobarbital (300 mg/kg) followed by cervical dislocation. Female mice were treated with fulvestrant (Sigma-Aldrich I4409) (150 mg/kg, i.p.) or its solvent (95% corn oil, 5% DMSO) 24 hours prior to acute UV exposure.

Chronic UV exposure: Females and males B6.129-SKH1-Hr*^hr^* aged 10-14 weeks old were exposed on their backs three times per week with a dose of 70mJ/cm2 delivered approximately in 1min and 45sec for 21 weeks. Each mouse received a total number of 74 UV doses and were then euthanized by lethal intraperitoneal injection of pentobarbital (300 mg/kg) followed by cervical dislocation. Mice were euthanized before the tumor reached 1cm^3^. Tumors size, appearance and number were monitored twice a week. Control mice were sham manipulated. Tumors, as well as proximal (1cm away from tumor) and distal (2cm away from tumor) skin from tumors were collected.

Multiple UV exposure for mutations detection: Females and males B6.129-SKH1-Hr*^h^*aged 10-14 weeks old were exposed on their backs three times per week with a dose of 70mJ/cm2 delivered approximately in 1min and 45sec for 2 weeks, 8 weeks or 12 weeks and aged for 18 weeks before being euthanized.

### Tumor grading and epidermal atypia grading

Mouse skin tumors derived from chronic or acute UV exposure were fixed in 4 % paraformaldehyde for 24 hours and then embedded in paraffin. Tissue sections (4µm) were stained with hematoxylin and eosin. Histological analysis of actinic keratosis and tumor classification were performed blindly by a dermatopathologist. SCCs were classified according to the Broders’ classification based on the degree of SCC keratinization and of keratinocyte differentiation (Broders AC., 1921). The classification includes SCC Grade I (>75% well-differentiated keratinocytes), SCC Grade II (25%-75% well-differentiated keratinocytes), SCC Grade III (<25% well-differentiated keratinocytes). Actinic keratosis was histologically defined and classified as grade I, II or III based on the degree of cytological atypia of epidermal keratinocytes and involvement of adnexal structures according to the criteria proposed by Rowert-Huber et al. in 2007 ^62^.

Epidermal atypia was graded based on cytological features (nuclear size, shape, and texture, as well as nucleolar prominence) and architecture changes (acanthosis, hyperkeratosis, change in epidermal thickness, and disorganization of cell alignment) ^63–67^. The degree of atypia was classified as follows: Grade 0 – no atypia, Grade 1 - discrete atypia, Grade 2 - intermediate atypia, Grade 3 - severe atypia. Histological assessment was performed blindly by a dermatopathologist.

### Whole-mount preparation of mice dorsal epidermis

The dorsal skin was cut into 24 pieces of 4×5 mm per mice and incubated in PBS containing 5mM EDTA at 37°C for 2.5 hours. Samples were transferred into PBS and the epidermis was carefully scraped off using curved scalpel while holding one corner of the skin with forceps. The epidermal wholemounts were fixed in 4% paraformaldehyde in PBS for 30 minutes and then stored in PBS at 4°C.

### Ultra-deep targeted sequencing Epidermal whole mounts

Wholemount epidermis was micro-dissected into a gridded array of 2mm^2^ samples. DNA was extracted using QIAGEN DNA micro kit (QIAGEN) by digesting overnight and following manufacturer’s instructions. DNA was eluted using pre-warmed AE buffer and was passed through the column a total of three times. DNA was extracted in the same manner from liver samples from the same animal to act as a germline control.

200ng of genomic DNA was fragmented to give an average size distribution of approx 150bp (LE220, Covaris Inc). Samples were purified, libraries prepared (NEBNext Ultra II DNA Library prep Kit, New England Biolabs), and index tags applied (Sanger 168 tag set). Index tagged samples were amplified (6 cycles of PCR, KAPA HiFi kit, KAPA Biosystems), quantified (Accuclear dsDNA Quantitation Solution, Biotium), then pooled together in an equimolar fashion. 500ng of pooled material was taken forward for hybridization, capture and enrichment (SureSelect Target enrichment system, Agilent technologies). A target bait panel of 74 genes was used. Genes were selected to cover those frequently mutated in squamous cell carcinoma.

The gene sequenced were:

*Aff3, Ajuba, Arid1a, Arid2, Arid5b, Atm, Atp2a2, Bcl11b, Braf, Cacna1d, Card11, Casp8, Ccnd1, Cdkn2a, Cobll1, Crebbp, Ctcf., Ctnnb1, Dclk1, Dclre1a, Dnmt3a, Ddr2, Egfr, Eif2d, Ep300, Erbb2, Erbb3, Erbb4, Ezh2, Fat1, Fat2, Fat3, Fat4, Fbxw7, Fbxo21, Fgfr3, Flt3, Grin2a, Hras, Kdm6a, Kdr, Kit, Kmt2c, Kmt2d, Kras, Lrp1b, Mtor, Nf1, Nf2, Notch1, Notch2, Notch3, Notch4, Nras, Pik3ca, Ptch1, Pten, Rb1, Ros1, Smad4, Smarca4, Smo, Sox2, Stat5b, Tert, Tet2, Tgfbr1, Tgfbr2, Trp53, Tsc1, Vhl, Zfp750, Nrf2, Keap1*.

Post-clean up samples were normalized to approximately 6nM and submitted to cluster formation for sequencing on Novaseq 6000 (Illumina) to generate 100bp paired-end reads.

BAM files were mapped to the GRCh37d5 reference genome using BWA-mem (version 0.7.17)^68^ Duplicate reads were marked using SAMtools (v1.11)^69^. Depth of coverage was also calculated using SAMTools to exclude reads which were: unmapped, not in the primary alignment, failing platform/vendor quality checks or were PCR/Optical duplicates. BEDTools (version 2.23.0) coverage program was then used to calculate the depth of coverage per base across samples^70^.

Mutation variant calling was made using the deepSNV R package (also commonly referred to as ShearwaterML), version 1.21.3, available at https://github.com/gerstung-lab/deepSNV, used in conjunction with R version 3.3.0 (2016-05-03)^71^. Variants were annotated using VAGrENT^72^. Mutations called by ShearwaterML were filtered using the following criteria:

- Positions of called SNVs must have a coverage of at least 100 reads.
- Germline variants called from the same individual were removed from the list of called variants.
- Adjustment for FDR and mutations demanded support from at least one read from both strands for the mutations identified.
- Pairs of SNVs on adjacent nucleotides within the same sample are merged into a dinucleotide variant if at least 90% of the mapped DNA reads containing at least one of the SNV pair, contained both SNVs.
- Identical mutations found in multiple contiguous tissue biopsies are merged and considered as a single clone to prevent duplicate clone counting.

ShearwaterML was run with a normal panel of approximately 45k reads.

COSMIC signature profiles (v3.2) (https://cancer.sanger.ac.uk/signatures) were used for Single Base Substitutions (SBS), and Double base Substitutions (DBS) signature classification using the SigProfiler packages as follows: MatrixGenerator (v1.2.12), Extractor (v1.1.12), Assignment (v0.0.13), Plotting (1.2.2). Frequency of mutations within each trinucleotide context was calculated using SigProfiler within the SBS288 context.

Comparison of differential selection was calculated as described ^73^.

### Human skin biopsies

Patient-derived formalin-fixed paraffin-embedded blocks with actinic keratosis and cutaneous SCC were obtained from the VITA certified Dermatology Biobank (CHUV_2103_12) of the Lausanne University Hospital (CHUV), Switzerland. Informed consent was obtained from all enrolled patients.

### *Ex vivo* human skin culture

The tissue sources were registered within the department biobank validated by the Operational Committee of Biobanks and Official Registers of the Lausanne University Hospital CHUV (Biobank N#BB_029_DAL). Skin explants were obtained from women adult healthy patients undergoing abdominoplasty (phototype 1 or 2, Fitzpatrick scale). The study was approved by the local ethics committee. Written informed consent was obtained from all subjects. Immediately post-excision, the human skin was washed in sterile phosphate buffered saline (PBS) and subcutaneous fat was removed with a scalpel. Next, the skin was cut into 1×1 cm square. Skin pieces were placed into 6cm dishes containing DMEM semi-solid medium (DMEM /F12 with Glutamax-1 (Gibco 31331-028) containing 10% of FBS, 1% of P/S and 1% of multivitamins and 2.5% of agarose). The epidermis was exposed to air whereas the dermis was embedded in the semi-solid medium. Pieces of skin were incubated at 37°C in a humidified atmosphere containing 5% CO2, and the medium was changed every three days. To mimic human UV exposure, human skin explants were exposed to 10% of UVA (365nm) and 90% of UVB (312nm) at 3 different doses: 240mJ/cm^2^ (UVA: 24mJ/cm^2^; UVB: 216mJ/cm^2^), 480mJcm^2^ (UVA: 48mJ/cm^2^; UVB: 432mJ/cm^2^) or 720mJ/cm^2^ (UVA: 77mJ/cm^2^; UVB: 648mJ/cm^2^). Immediately after UV exposure skin was placed in fresh medium and incubator. Control skin not exposed to UV was sham manipulated.

### Immunofluorescence staining

Tumors and skin were fixed in 4% paraformaldehyde solution for 24h, then PBS washed, and paraffin embedded. Staining was performed on 4µm skin sections.

Antigen retrieval using Citrate buffer 0.01M pH6 was performed. Sections were blocked in NGS 5% for 45-90 minutes, incubated with the following antibodies : anti-CPD (Cosmo Bio, NMDND004, 1/1000), anti-γH2AX (EMD Millipore, 05-636-25UG, 1/500), anti-Ki67 (Invitrogen, 41-5698-80, 1/60), anti-Keratine10 (Progen, GP-K10, 1/100), anti-Keratine14 (Covance, PRB155P, 1/500) anti-cleaved caspase-3 (Cell Signaling Technology, 9664, 1/100), anti-CDKN3 (Abcam, ab175393, 1/500) in NGS 5% O/N at 4°C; washed in PBS; incubated with a fluorescent antibody (Molecular Probes, 1/500 in NSG 5%) for 30 minutes at room temperature: goat anti-mouse A568 (for CPD and γH2AX), goat anti-rat A568 (for KI67), Goat anti-guinea pig A488 (for K10), goat anti-rabbit A568 (for K14 and CDKN3), goat anti-rabbit A568 (for cleaved caspase-3), washed in PBS, counterstained with DAPI, and embedded in MOWIOL. Pictures were taken with a motorized AxioImager M1 microscope and the AxioVision software (Carl Zeiss), using a magnification of x10, ×20 or x40. After defining epidermis region, automatic cell detection of staining was performed using Qupath software ^74^ in a blinded way.

### Epidermis-dermis separation for RNA extraction

Epidermis and dermis were prepared from whole dorsal skin of mice according to the protocol developed by Clemmensen *et al* ^75^. After dorsal skin harvesting, the skin was cut into thin strips (2 mm wide) and immediately incubated for 20 min at RT in ammonium thiocyanate (3.8% in 1XPBS). Before mechanical separation with forceps and scalpel, skin was washed once with 4°C PBS.

### RNA isolation and RT-qPCR analysis

Total RNA was isolated from human and mouse epidermis using TRIzol LS (Life technologies, Thermo Fisher) and purified with the RNeasy kit (Qiagen) according to the manufacturer’s instructions. RNA quality was verified by microfluidic (Agilent 2100 Bioanalyzer) and concentration determined with a NanoDrop spectrophotometer (Wilmington). Total RNA (500ng – 1ug) was reverse transcribed using an RT Go script Kit (Promega) according to the manufacturer’s instructions. cDNA quantifications were performed with GoTaq qPCR Master Mix (Promega). No template control and no reverse transcriptase enzyme sample were used as negative controls. Gene expression was normalized to the mean of two housekeeping gene expression (*Eef1a1* and *Rpl27* genes). Primer sequences are accessible in the supplemental Table 1.

### RNA-seq gene expression studies

RNA-seq experiment was conducted at the Lausanne Genomic Technologies Facility (Lausanne, Switzerland) according to an in-house pipeline. Briefly, mouse RNA samples were extracted from four epidermal samples of each sex, in control epidermis and 24h after single UV exposure (120mJ/cm^2^). RNA quality was assessed on a Fragment Analyzer (Agilent Technologies) and all RNAs had a RQN from 8 to 10. RNA-seq libraries were prepared from 500 ng of total RNA with the Illumina TruSeq Stranded mRNA reagents (Illumina) using a unique dual indexing strategy and following the official protocol automated on a Sciclone liquid handling robot (PerkinElmer). Libraries were quantified by a fluorimetric method (QubIT, Life Technologies) and their quality assessed on a Fragment Analyzer (Agilent Technologies). Sequencing was performed on an Illumina NovaSeq 6000 for 100 cycles single read. Sequencing data were demultiplexed using the bcl2fastq2 Conversion Software (version 2.20, Illumina).

#### RNA-seq data processing

Data cleaning: Purity-filtered reads were adapted and quality trimmed with Cutadapt (v. 1.8, Martin 2011). Reads matching to ribosomal RNA sequences were removed with fastq_screen (v. 0.11.1). Remaining reads were further filtered for low complexity with reaper (v. 15-065).

Reads were aligned against the GRCh38.102 genome using STAR (v. 2.5.3a). The number of read counts per gene locus was summarized with htseq-count (v. 0.9.1) using GRCh38.102 gene annotation. Quality of the RNA-seq data alignment was assessed using RSeQC (v. 2.3.7).

Statistical analysis was performed in R (R version 4.1.0). Genes with low counts were filtered out according to the following rule: at least 1 sample had to have at least 1 cpm (1 count per million) reads in order to keep the gene in the data set. Library sizes were then scaled using TMM normalization. Subsequently, the normalized counts were transformed to cpm values and a log2 transformation was applied by means of the function cpm with the parameter setting prior.counts = 1 (EdgeR v 3.34.1).

#### Differential expression

Differential expressions were computed with the R Bioconductor package limma by fitting data to a linear model. The approach limma-trend was used. Results from contrasts of interest and interactions were extracted. Moderated F-tests were performed for groups of contrasts and groups of interactions. The resulting p-values were adjusted for multiple testing by the Benjamini-Hochberg method, which controls for the false discovery rate (FDR). This adjustment was performed for each F-test. A post-hoc test was performed, using the function decide Tests with parameter method=nestedF.

Functional annotation clustering of GO Biological Process was performed using DAVID Bioinformatics resource system ^76^.

### Publicly available datasets

Public datasets were found on the Gene Expression Omnibus data repository, two independent gene expression datasets were used: GSE66412 and GSE84293. GSE66412 access available in August 2015, compared the RNAseq whole transcriptome of Normal human epithelial keratinocytes (NHEKs) and Squamous cell carcinoma cell lines (SCC). GSE84293 access available on February 2017 through next generation sequencing compared transcriptome of normal skin (NS, peritumoral), actinic keratosis (AK) and cutaneous SCC. Values for CDKN3 and CDC25C gene expression from RNA sequencing data were calculated using GREIN ^77^.

### Kaplan-Meier plotter database analysis

The correlation between CDKN3 expression and CDC25C expression and survival in head and neck squamous cells carcinoma patients (n=500) was analyzed using Kaplan-Meier plotter (http://kmplot.com/analysis), the hazard ratio (HR) with 95% confidence intervals and log-rank P-value was also determined ^78^.

### Statistical analysis

To compare two independent groups, we used the student *t-*test; to compare one independent variable in more than two groups, we used the one-way ANOVA with Tukey’s post hoc test; to compare two independent variables in more than two groups, we used two-way ANOVA with Sidak’s post hoc test. Statistics on the survival curves were performed using a log-rank t-test. All statistical analyses were performed using Prism GraphPad (v7)

### Data availability

The sequencing dataset generated during this study is available at the European nucleotide Archive (ENA) with the following dataset accession number: ERP159842 (https://www.ebi.ac.uk/ena/browser/view/PRJEB75253). The data generated during this study have been deposited in the GEO database under accession code GSE266221 (https://www.ncbi.nlm.nih.gov/geo/q).

## Acknowledgements

We thank the Animal facility (Center of Integrative Genomics, University of Lausanne, Switzerland) for expert technical assistance and the Genomic Technologies Facility (University of Lausanne, Switzerland) for performing RNA-seq analyses. We thank Fabienne Lammers (Center for Integrative Genomics, University of Lausanne, Switzerland) for mouse genotyping and Nathalie Hirt (Lausanne University Hospital, University of Lausanne, Switzerland) for valuable support in getting human skin samples from the Lausanne University Hospital biobank. We are grateful to Prof. Hon. Béatrice Desvergne (University of Lausanne, Switzerland) and Hervé Guillou, PhD, Laurence Payrastre, PhD and Sandrine Ellero-Simatos, PhD (Toxalim, INRAE, ENVT, INP-PURPAN, UMR 1331, UPS, University of Toulouse, France) and Alexandra Montagner, PhD (I2MC, UMR1297, INSERM/UPS), Dorian Ziegler, PhD and Sarah Geller, PhD (Center of Integrative Genomics, University of Lausanne, Switzerland) for constructive scientific discussion.

This work was supported by the Swiss National Science Foundation (IZLIZ3_200253/1 to EG), the Foundation Recherche Cancer ISREC (CCP 10-3224-9 to EG), and the Etat de Vaud (University of Lausanne; SKINTEGRITY.CH collaborative research to EG and LM).

## Authors contributions

C.L. designed and supervised all experiments, conducted experiments, analyzed and interpreted data, prepared figures and wrote the manuscript. C.W. bred the mice, conducted experiments, analyzed data, and contributed to manuscript review and editing. C.R. developed immunological staining and provided all the histological data. J.C.F. performed DNA sequencing of mouse epidermis, analyzed and interpreted the sequencing data, contributed to experimental design and to manuscript writing, review and editing. Y-C.T. contributed to “material and method” section writing, compiled tumor analysis, selected representative photos, and contributed to tumor section analyses and grading. S.C. contributed to in vivo experiments and to manuscript review and editing. J.M. collected and provided human skin explants and contributed to manuscript review and editing. Y-T.C. coordinated mouse tumor slide sorting, renaming and scanning, setup scanning conditions, contributed to “material and method” section writing, and to manuscript review and editing. C.I. identified and collected patient AK and SCC samples, managed the contacts with the hospital biobank, and double-checked all diagnoses. P.H.J. contributed to DNA sequencing experimental design, to interpretation of the data, and to manuscript review and editing. E.G. graded mouse epidermis atypia and tumor samples, confirmed diagnoses of patient samples, selected representative photos, contributed to the “material and method” section writing, to interpretation of data and to manuscript review and editing. P.J. contributed to the design of the manuscript, contributed to analyses and interpretation of data, and to manuscript review and editing. L.M. contributed to the design and supervised the study, to the interpretation of data and wrote the manuscript.

