## Supplemental Figures for "Estrogen Receptors/E2F1/CDKN3 Axis Protects from UV-induced Skin Cancers in Females"

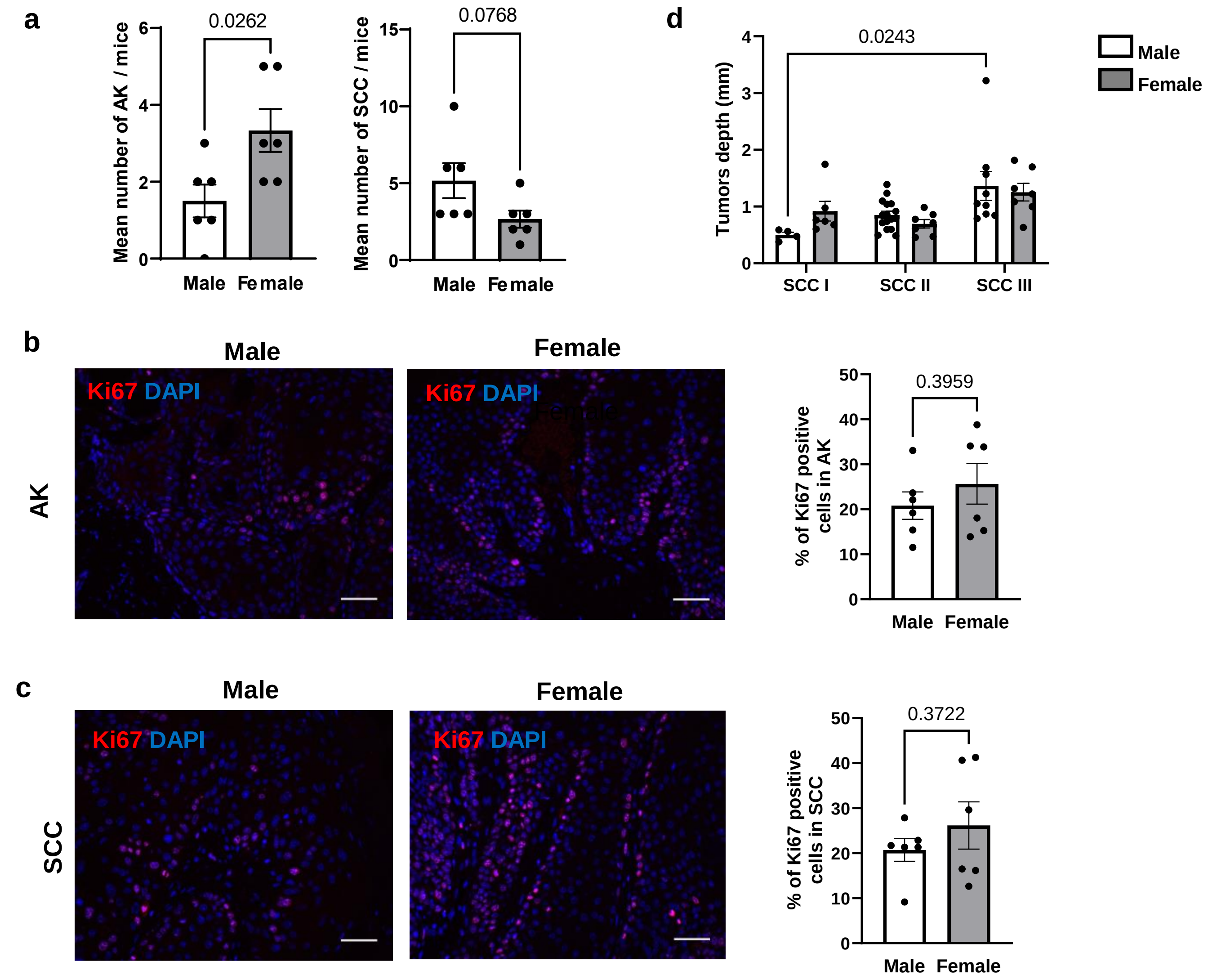


**Supplementary Fig. S1. Characterization of dorsal lesions following chronic UV exposure in males and in females.**

**a.** Mean number of actinic keratosis (AK) and squamous cell carcinoma (SCC) lesions collected per mice. *n= 6 for each male and female mice group*, mean ± SEM, Unpaired *t*-test.

**b.** Tumor depth from different stages SCC coming from male (white) or female (grey) mice chronically exposed to UV. Each dot represents a tumor, *n=4-16 per group*. Mean± SEM, two-way ANOVA with Sidak's post hoc test.

**c-d. Left:** Ki67 (red) immunofluorescence in AK (panel c) or SCC (panel d) lesions from male and female mice chronically exposed to UV. DAPI was used as counterstaining (blue). Scale bars: 50 μm. **Right:** Quantification of the percentage of Ki67 positive keratinocytes. *n(fields)= 3 per mice*, Mean± SEM, Unpaired *t*-test.

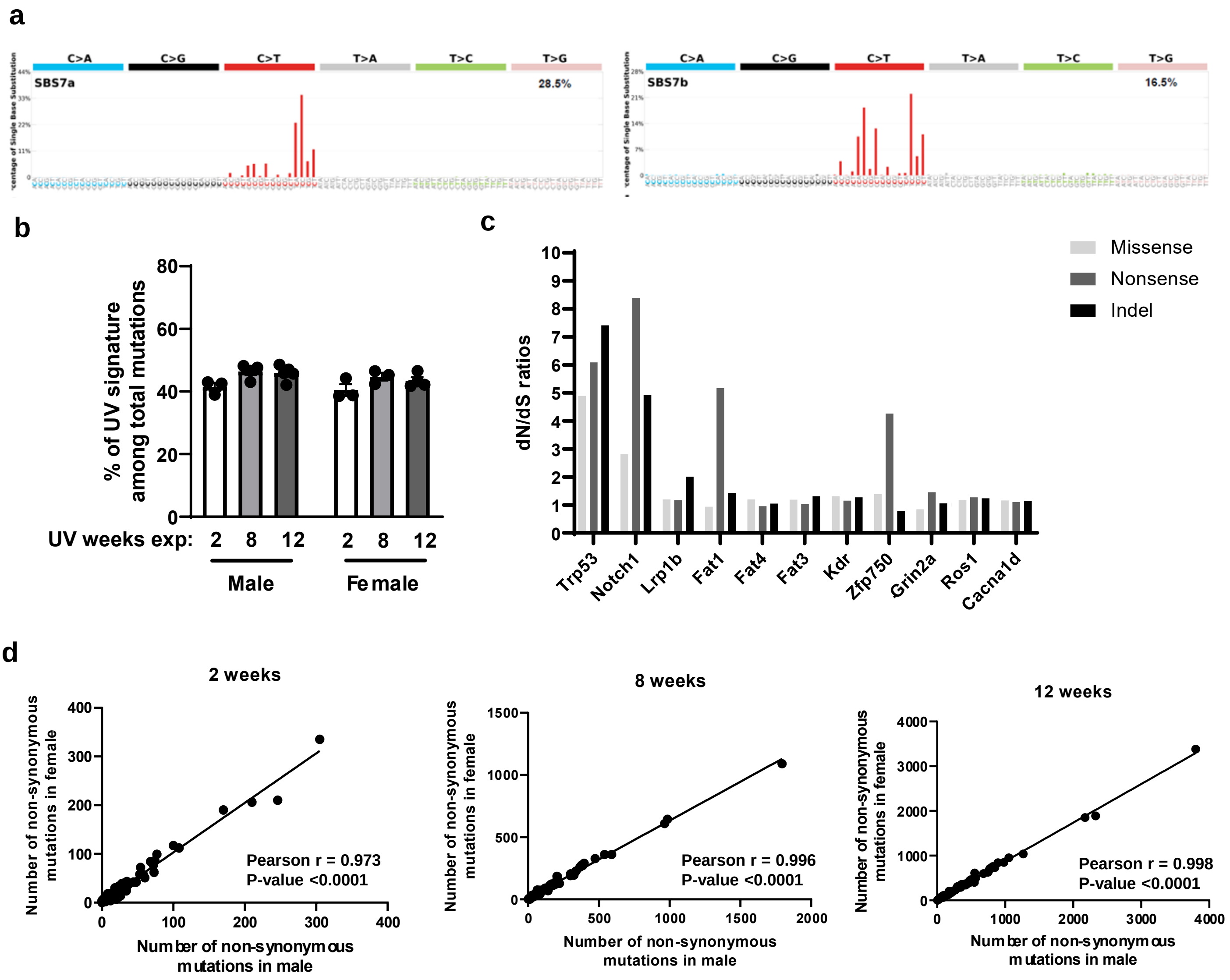


**Supplementary Fig. S2. Characterization of mutation types based on sex.**

**a.** UV mutational signatures (SBS7a and SBS7b) on epidermis from mice exposed to UV for 8 weeks and aged for 14 weeks.

**b.** Percentage of mutations attributed to UV derived signatures SBS7a and SBS7b in male and female dorsal epidermis collected from mice exposed to UV (70mJ/cm²) for 2 weeks, 8 weeks or 12 weeks, n= 3 to 5 mice, mean ± SEM.

**c.** Genes under positive selection across the entire data set using dN/dScv. Only genes with q<0.01 are shown.

**d.** Differential selection between male and female mice at different time points shows no significant difference.

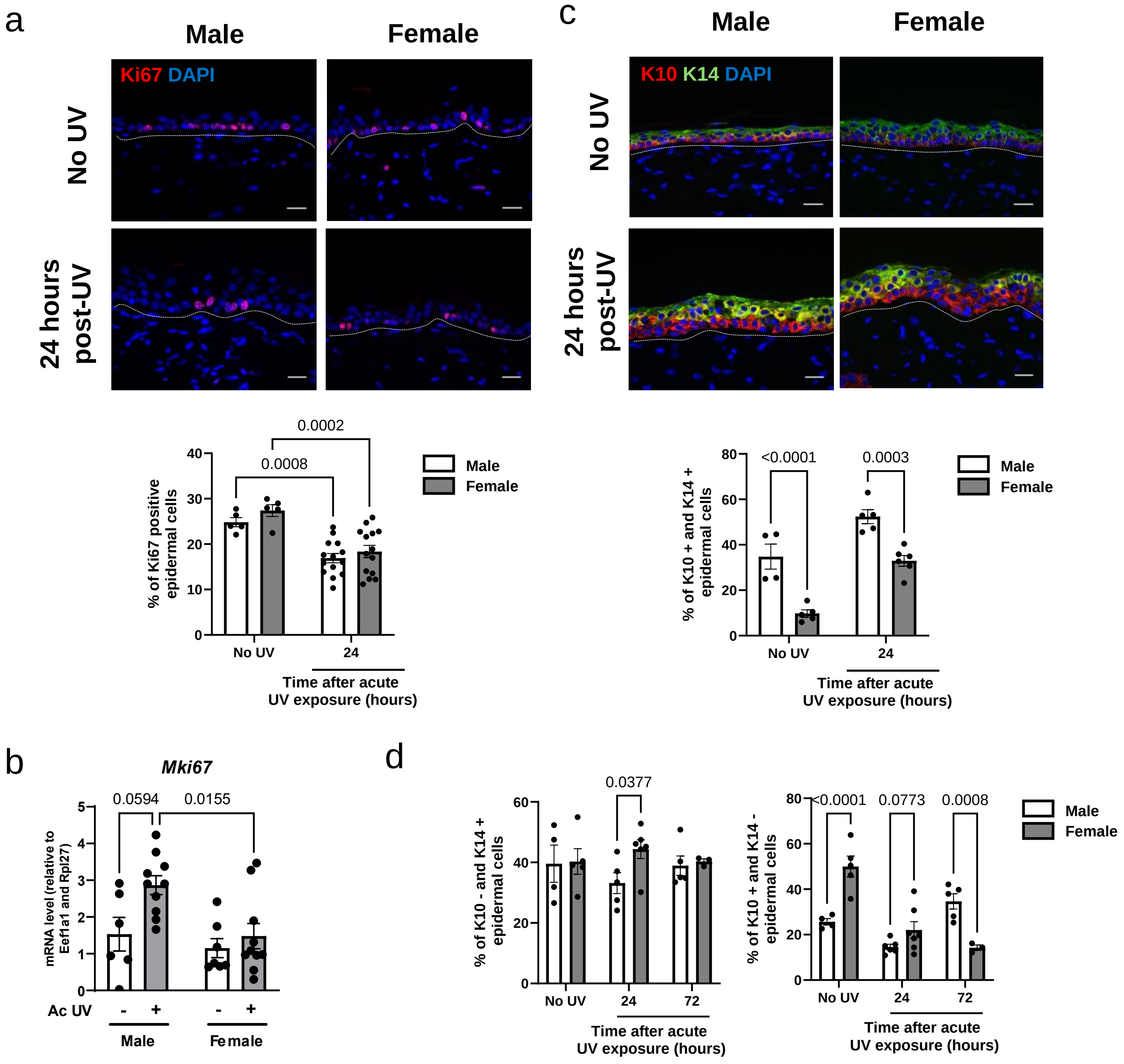


**Supplementary Fig. S3. Epidermal proliferation and differentiation following acute UV exposure in males and females.**

**a. Top:** Ki67 (red) immunofluorescence in male and female dorsal epidermis collected 24 hours after a single dose of acute UV exposure (120mJ/cm²), compared to control skin (No UV). DAPI was used as counterstaining (blue). The dotted line separates the epidermis from the dermis. Scale bar: 20 µm. **Bottom:** Percentage of Ki67 positive keratinocytes. *n(fields)= 4 per mouse*, mean ± SEM, two-way ANOVA with Sidak's post hoc test.

**b. Top:** Keratin 14 (K14; red) and Keratin 10 (K10: green) immunofluorescences in male and female dorsal epidermis collected 24 hours after a single dose of acute UV exposure (120mJ/cm²), compared to control skin (No UV). DAPI was used as counterstaining (blue). The dotted line separates the epidermis from the dermis. Scale bar: 20 µm. **Bottom:** Percentage of K10 positive and K14 positive epidermal cells. *n(fields)= 4 per mouse*, mean ± SEM, two-way ANOVA with Sidak's post hoc test.

**c.** RT-qPCR analysis of Mki67 mRNA expression. *n= 5 to 10 mice*, mean ± SEM, two-way ANOVA with Sidak's post hoc test.

**d.** Percentage of K10 negative and K14 positive (left), and K10 positive and K14 negative (left) epidermal cells. *n(fields)=4 per mouse, n= 3 to 6 mice*, mean ± SEM, two-way ANOVA with Sidak's post hoc test.

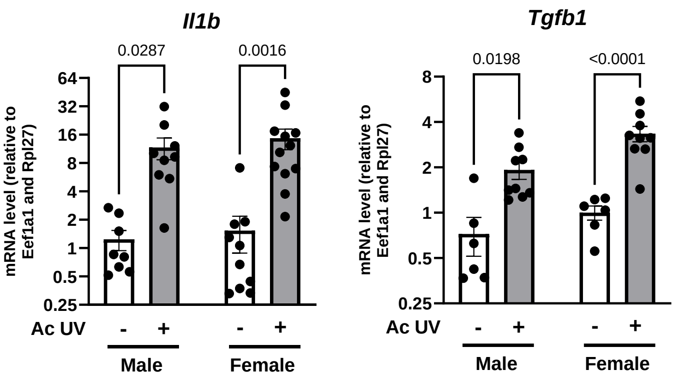


**Supplementary Fig. S4. Cytokine production in male and female mice following acute UV exposure.**

Quantification of the relative gene expression of *Il1b* and *Tgfb1* at mRNA level by RT-qPCR in male and female mice epidermal samples. *n=7-12 mice per sex*, Mean ± SEM, two-way ANOVA with Sidak's post hoc test.

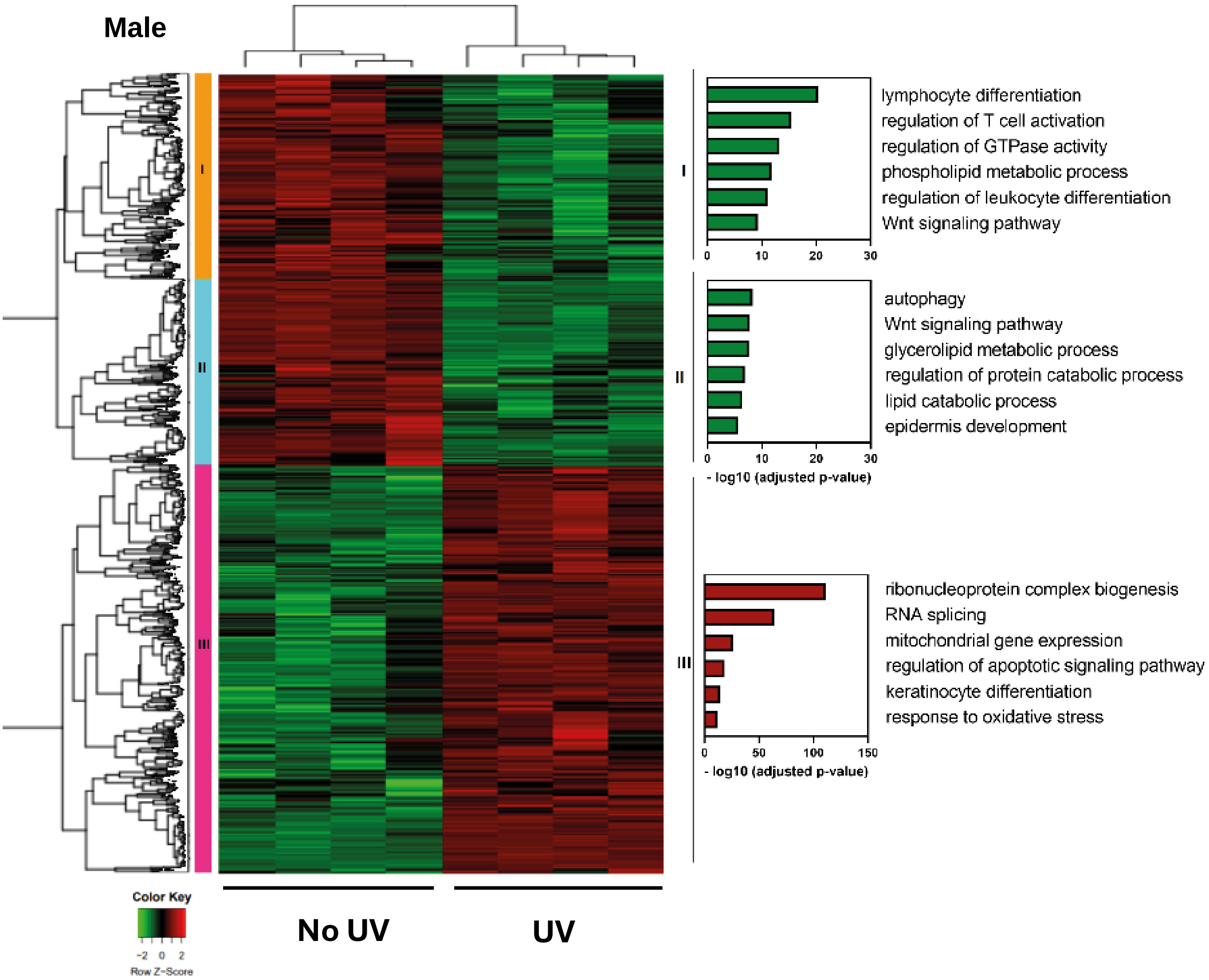


**Supplementary Fig. S5. Heatmap showing differentially expressed genes in male epidermis following UV exposure (UV) *vs. n*on-UV exposure (No UV).**

Differentially expressed genes clustering heat map for RNA-seq data showing the log2 transformed expression values of individual samples in no UV-exposed and UV-exposed (single dose; 120mJ/cm$^2$) epidermal samples in male mice. Each row represents one gene. Log2 expression values for each single gene are resized to row z-score scale (from −2, the lowest expression to +2, the highest expression for single gene), colors represent gene expression changes. Red indicates upregulation of gene expression and green indicates downregulation of expression. Right: Significantly enriched pathways in each cluster as determined by Reactome Pathway Enrichment analysis. *n= 4 mice per sex*

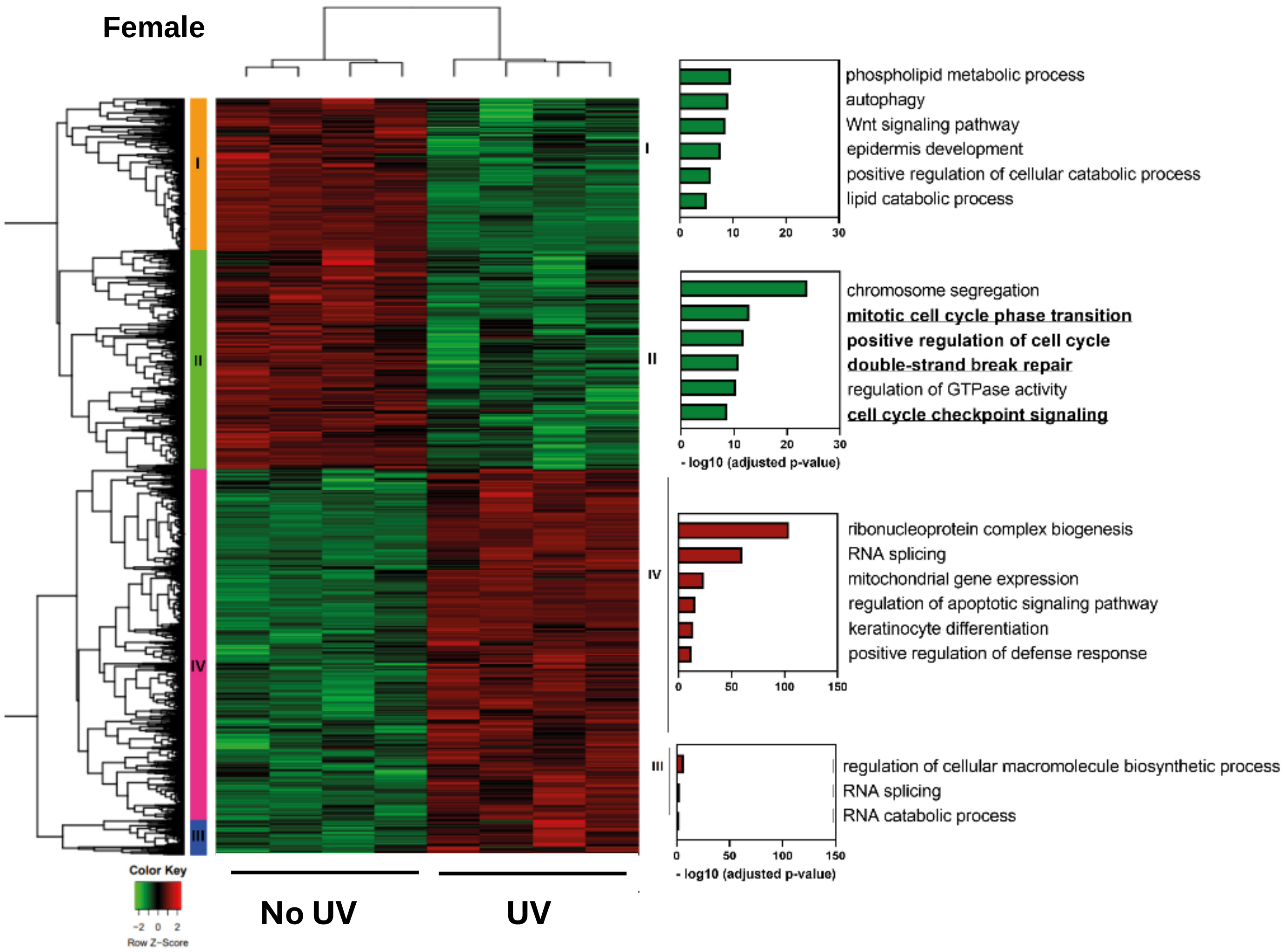


**Supplementary Fig. S6. Heatmap showing differentially expressed genes in female epidermis following UV exposure (UV) *vs.* non-UV exposure (No UV).**
Differentially expressed genes clustering heat map for RNA-seq data showing the log2 transformed expression values of individual samples in no UV-exposed and UV-exposed (single dose; 120mJ/cm$^2$) epidermal samples in female mice. Each row represents one gene. Log2 expression values for each single gene are resized to row z-score scale (from −2, the lowest expression to +2, the highest expression for single gene), colors represent gene expression changes. Red indicates upregulation of gene expression and green indicates downregulation of expression. Right: Significantly enriched pathways in each cluster as determined by Reactome Pathway Enrichment analysis. . *n= 4 mice per sex*

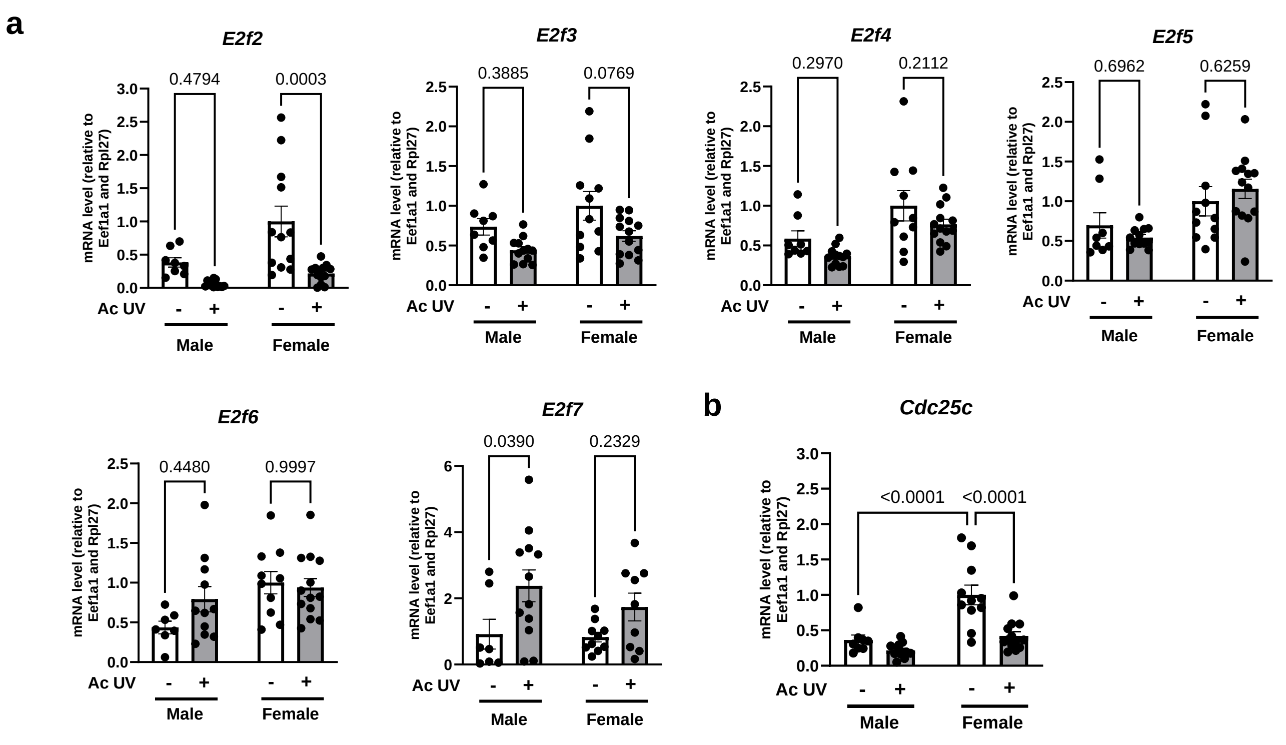


**Supplementary Fig. S7. Relative gene expression in male and female epidermis following acute UV exposure.**

**a.** Quantification of the relative gene expression of *E2f2, E2F3, E2f4, E2f5, E2f6 and E2f7* at mRNA level by RT-qPCR in male and female mice epidermal samples. *n=8-14 mice per sex*, Mean ± SEM, two-way ANOVA with Sidak's post hoc test.

**b.** Quantification of the relative gene expression of *E2f2 and Cdc25c* at mRNA level by RT-qPCR in male and female mice epidermal samples. *n=8-14 mice per sex*, Mean ± SEM, two-way ANOVA with Sidak's post hoc test.

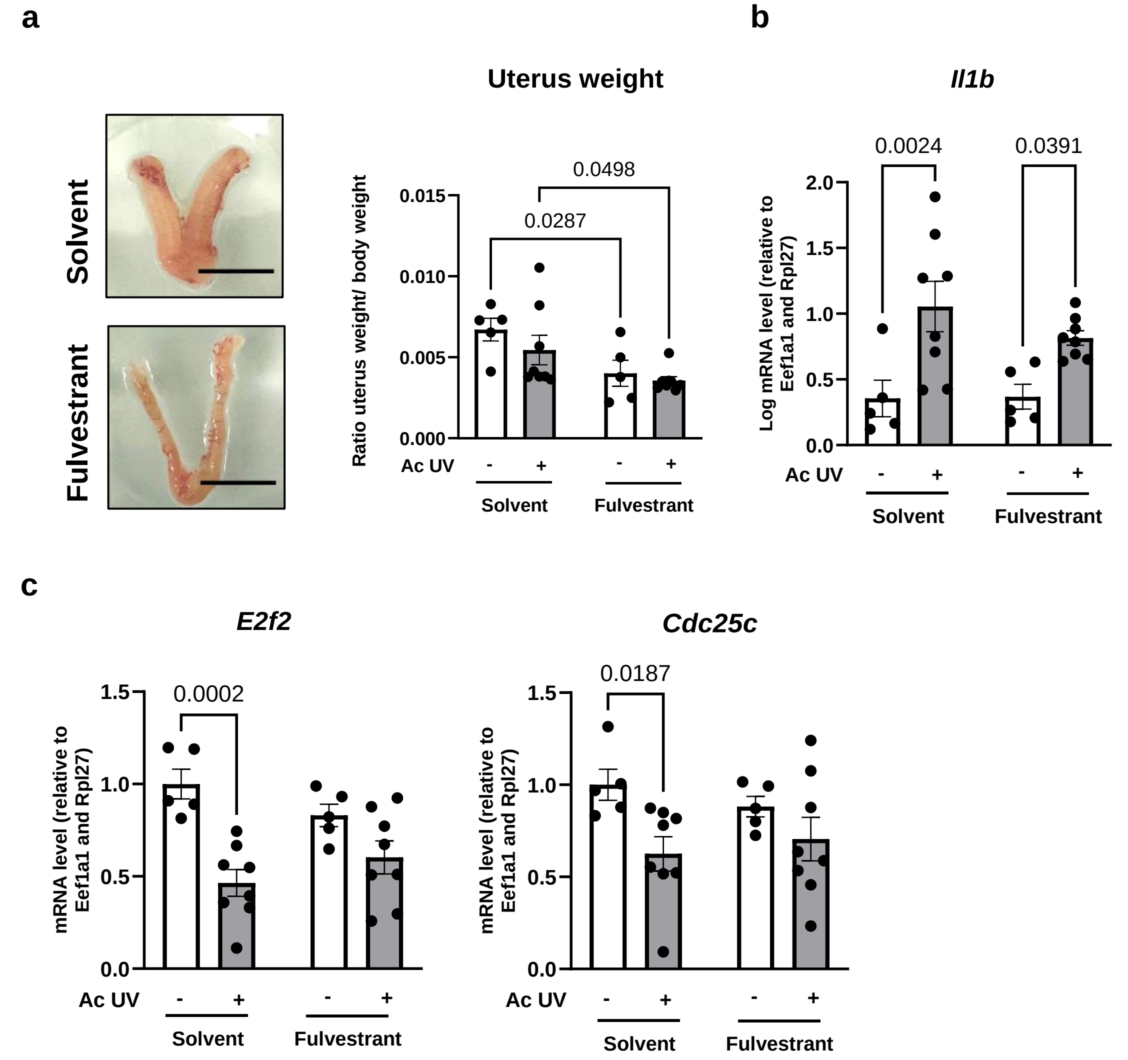


**Supplementary Fig. S8. Effect of fulvestrant treatment on female mice exposed to acute UV.**

**a. Left:** Representative image of the female uterus following treatment with either solvent or fulvestrant. **Right:** Relative uterus weight after treatment with fulvestrant (150 mg/kg) for 48 hours, or with solvent (control*). n = 5-8 mice*, Mean ± SEM, two-way ANOVA with Sidak's post hoc test.

**b.** Quantification of the relative gene expression of *Il1b* at mRNA level by RT-qPCR in female mice epidermal samples. *n=5-8 mice*, Mean ± SEM, two-way ANOVA with Sidak's post hoc test

**c.** Quantification of the relative gene expression of *E2f2* and *Cdc25c* at mRNA level by RT-qPCR in female mice epidermal samples. *n=5-8 mice*, Mean ± SEM, two-way ANOVA with Sidak's post hoc test

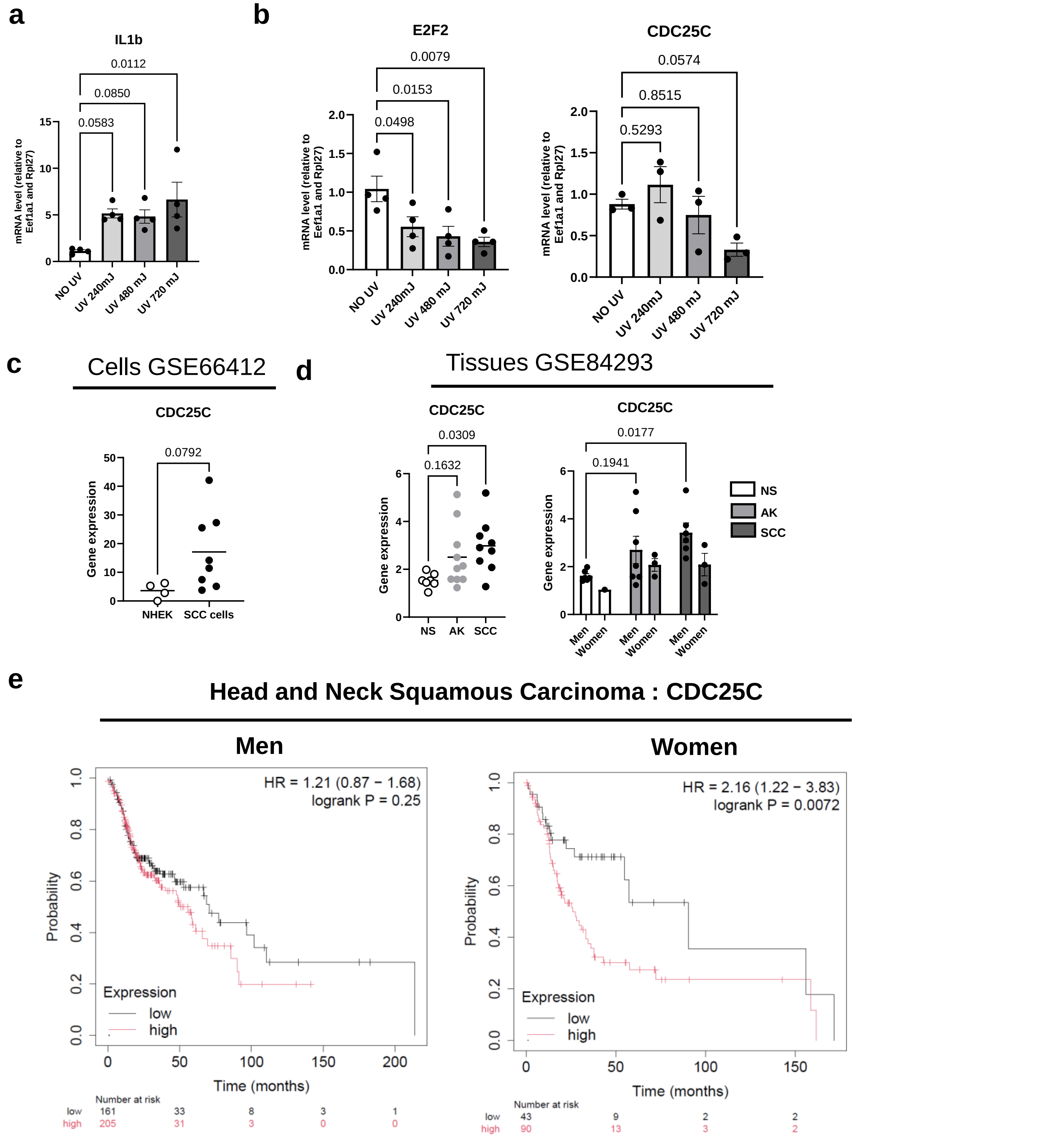


**Supplementary Fig. S9**. **Downregulation of CDC25C expression in the skin of healthy women and in squamous cell carcinoma.**

**a-b** Relative mRNA expression of *IL1b, E2F2* and *CDC25C* in *ex vivo* skin explant cultures from women subjects collected 24 hours after exposure to a single dose of UV with increasing intensities (240, 480 and 720 mJ/cm2) compared to non-UV-exposed control explants, quantify by RT-qPCR. *N=4 subjects*. Mean ± SEM, two-way ANOVA with Tukey's post hoc test.

**c.** Expression of *CDC25C* mRNA expression in normal human epithelial keratinocytes (NHEK) and human Squamous Cell Carcinoma (SCC). Data from available public datasets (GSE66412). NHEK: *n= 4*, SCC: *n= 8*. Mean ± SEM, t- student test.

**d.** Expression of *CDC25C* mRNA expression in men and women human Normal Skin (NS), Actinic Keratosis (AK) and Squamous Cell Carcinoma (SCC) lesions. Data from available public datasets (GSE84293). NS: *n= 7 (NS; 6 men and 1 woman), n= 10 (AK; 7 men and 3 women), n=9 (SCC; 5 men and 3 women).* The two right graphs represent gene expression with sex separation. Mean ± SEM, one-way ANOVA with Tukey's post hoc test for non sex separated graph, two-way ANOVA with Tukey's post hoc test for sex-separated graphs.

**e.** Kaplan-Meier survival curves for men (left) or women (right) patients with Head and Neck Squamous Cell Carcinoma (HNSC) based on CDC25C gene expression by Kaplan Meier plotter.
